# Remodeling of mRNA by eIF4F in human translation initiation

**DOI:** 10.64898/2026.06.19.733470

**Authors:** Carlos Alvarado, Christopher P. Lapointe, Jinfan Wang, Masaaki Sokabe, Rosslyn Grosely, Ajinkya A. Dhepe, Crystal I. Stackhouse, Michael Z. Palo, Adam Mamot, Jacek Jemielity, Christopher S. Fraser, Joseph D. Puglisi

## Abstract

For the ribosome to load onto an mRNA during the early steps of translation initiation, the mRNA must be activated by the eIF4F complex. The mechanism of this activation step has remained elusive. Here we employ multi-perspective real-time single-molecule assays to observe directly mRNA-eIF4F binding near the 5′ end, mRNA conformational remodeling, and 40S ribosomal subunit loading. eIFs 4E, 4G, and 4B play distinct roles in promoting eIF4F association and stabilizing eIF4A binding. Binding of eIF4F is the rate-limiting step in mRNA activation: once bound, mRNA conformation is rapidly extended in an ATP-dependent manner. The mRNA extended state is the necessary substrate for 43S PIC loading and perturbations to extension delay loading. Features of the mRNA, such as the 7-methylguanosine cap at the 5′ end and secondary structures, modulate these steps and regulate ribosome loading. Our results establish a kinetic and mechanistic framework for the early steps in translation initiation.

## Introduction

Human translation initiation selects a messenger RNA (mRNA) template and determines the start site of protein synthesis, controlling the identity and amount of protein products^1–5^. A typical human cell can contain 10^5-^10^6^ mRNA molecules^6^, which compete for limited initiation factors and ribosomes. Decades of biochemical, biophysical, and genetic studies have defined the eukaryotic initiation factor (eIF) 4F complex as the set of proteins that bind to and activate an mRNA for 40S ribosomal loading near the 5′ 7-methylguanosine cap (5′ cap)^7–9^. Loaded 40S subunits, containing other eIFs and initiator tRNA then scan directionally to find the AUG start codon with subsequent factor rearrangement steps and assembly of an elongation-competent 80S ribosome^2–4,10^. Remodeling of mRNA conformation is a proposed sub-step in mRNA activation and likely driven by eIF4F^7,9^. How exactly an mRNA is remodeled and activated by eIF4F for the loading of a 40S ribosomal subunit is not known.

The eIF4F complex is composed of eIF4A, eIF4E, and eIF4G. This complex is scaffolded by eIF4G1 (hereafter referred to as eIF4G), which contains a binding site for eIF4E and two binding sites for eIF4A1 (hereafter referred to as eIF4A), alongside other binding sites for eIF3, poly(A)-binding protein and RNA^11–16^. Assembly of the multiprotein eIF4F complex can occur in the absence of mRNA, though it remains unclear in canonical initiation whether preformed eIF4F complexes associate to an mRNA or if the complex assembles on the mRNA. eIF4F is targeted to the 5′ end of an mRNA through interactions between eIF4E and the 5′ cap of an mRNA^17,18^. Thus, the cap serves as a docking site that orients the complex toward the end of an mRNA that is conducive for subsequent steps of initiation^1^. Recent studies in yeast demonstrated that the 5′ cap stabilizes eIF4F binding in contrast to off-pathway binding at interior sites of the mRNA^19^.

eIF4F remodels mRNA conformation at the 5′ end in preparation for 40S subunit loading. Remodeling is likely facilitated by the DEAD-box helicase and ATPase eIF4A, which unwinds mRNA secondary structures^20^. The helicase and ATPase activities of eIF4A are stimulated by both eIF4G and eIF4E but also by eIF4B^21–28^, an RNA-binding protein that transiently interacts with eIF4A on mRNA^23,29–31^, and is required for initiation^32,33^. Human mRNAs often contain complex secondary structures in the 5′ untranslated region (5′ UTR), but only single-stranded mRNA can be threaded through the mRNA binding channel on the 40S subunit^34^. Therefore, during mRNA activation, secondary structure must be remodeled prior to and during 40S subunit loading. However, studies have shown eIF4A on its own to be a relatively weak helicase^21,27,35,36^, unable to unwind complex mRNA secondary structures. To what extent eIF4A in conjunction with the other initiation factors can remodel mRNA structure and what the necessary remodeling is for ribosomal loading remains unclear.

After an mRNA has been activated by eIF4F, the 40S ribosomal subunit loads near the 5′ end^1,4^. The 40S subunit loads in complex with eIFs 1, 1A, 3, and the eIF2-GTP-Met-tRNA_i_ ternary complex (TC), as a 43S preinitiation complex (PIC)^2,3,37^. Loading is facilitated by an interaction network involving the mRNA, eIF4G, eIF3, and the 40S subunit^1,38,39^. eIF4A and 4B can also directly bind to the 40S subunit^40,41^, which could contribute to PIC loading. Upon loading onto the 5′ end of an mRNA, the PIC scans in a net 3ʹ direction until a start codon is recognized^2,42^. Conformational and compositional rearrangements of the PIC induced by eIFs 5 and 5B select the start codon and facilitate the binding of the 60S subunit^1,4,37,43,44^. After departure of eIFs, the elongation competent 80S initiation complex is positioned at the start codon.

Multiple signaling pathways regulate eIF4E availability, eIF4F complex phosphorylation and formation^8,9,45^. Dysregulation of these factors and related pathways occurs in various cancers, leading to overexpression of these proteins and an unregulated increase in translation^33,46–49^. Additionally, certain viruses target eIF4F by producing proteases that cleave eIF4G, reducing host mRNA translation while promoting viral RNA cap-independent translation pathways^9,50–53^. Recent studies have provided further insights into the timing of cap-recognition by eIF4F in yeast but not in humans^18,19^.

Here, we developed several single-molecule Förster resonance energy transfer (FRET) assays using our established *in vitro* reconstituted human translation initiation system^43^, to define mRNA activation by eIF4F. First, we monitor eIF4F binding kinetics proximal to the 5′ cap, using fluorescently labeled eIF4A within eIF4F as a reporter for the full complex. Then we tracked conformational remodeling of mRNA by eIF4A and cofactors in real-time, and finally correlated mRNA remodeling in the context of 43S PIC loading. We demonstrate that mRNA remodeling is the activation step and productive 43S PIC loading is contingent on eIF4F/eIF4B-mediated remodeling. These findings establish a mechanistic framework for the early steps in translation initiation.

## Results

### Dynamics of eIF4F binding proximal to the cap

To examine directly how eIF4F binds to the 5′ UTR adjacent to the cap, we established a single-molecule FRET assay using a four-color detection zero-mode waveguide (ZMW)^54^ platform to monitor eIF4A within eIF4F, as a reporter for the full complex (Fig. 1a). We fused a ybbR-tag^55^ to the N-terminus of eIF4A, which did not perturb RNA-dependent ATPase activity (Fig. S1a), and was fluorescently labeled via conjugation with a Cy5 dye. Full-length human β-globin mRNA was fluorescently labeled via Cy3 dye conjugation at the 5′ cap using an azido-functionalized cap analog (N_3_-m^7^GpppG) (Fig. S1b), which provides a unique handle for site-specific labeling while retaining the ability to support cap-dependent translation. Functionality of this analog was previously validated *in cellulo*^56^. The Cy3-cap-mRNA was tethered to an imaging surface within ZMWs, and the surface was excited directly by a 532-nm laser (Fig. 1a). For initial characterization of our signal, we added Cy5-labeled eIF4A (100 nM) and saturating levels of adenosine triphosphate (ATP) in real time. As predicted, we observed Cy5-eIF4A binding proximally to the Cy3-cap of mRNA as pulses of Cy3-to-Cy5 FRET (Fig. 1b). The mean elapsed time from solution mixing to when eIF4A bound – the mean eIF4A association time – was 455 ± 62 s (Fig. 1d). This value represents a time-censored lower bound, as it is very close to our observation window. The mean bound state lifetime before eIF4A departure was 10.8 ± 0.2 s (Fig. S1f). The rates and rate constants used to derive mean times are listed in Table S1.

**Figure 1.**
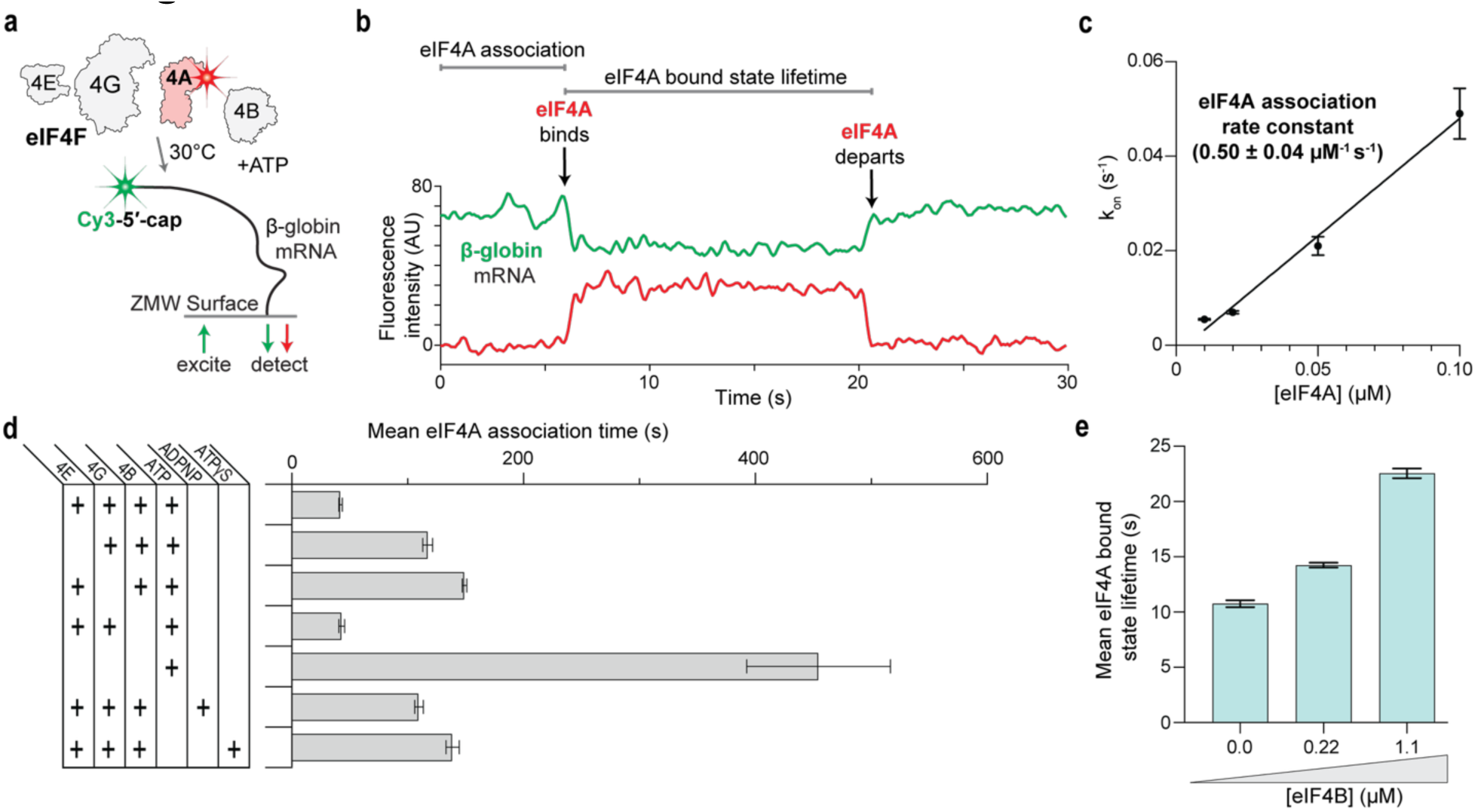
Dynamics of eIF4F binding proximal to the cap. **a.** Schematic of single-molecule 5′ end-to-eIF4A FRET assay where Cy5-labeled eIF4A (red) alongside eIF4E, eIF4G, eIF4B, and ATP was delivered to Cy3-5′-cap-labeled β-globin mRNA (green) tethered on a ZMW imaging surface. **b.** Example single-molecule data for the assay, FRET corresponds to the anti-correlated decrease in green signal and increase in red signal. The time taken for FRET to appear is the eIF4A association time, the lifetime of FRET is designated the eIF4A bound state lifetime. **c.** Plot of observed eIF4A association rate across different concentrations. Linear regression analysis (solid line) was used to derive the association rate constant. The error bars represent the 95% confidence interval (CI) of the observed association rate constants. From lowest to highest concentration: n = 146, 126, 131, and 121. **d.** Plot of mean eIF4A association time across various conditions. Error bars represent the derived 95% CI of the mean association times. From top to bottom: n = 115, 99, 202, 105, 196, 142, and 77. **e.** Plot of mean eIF4A bound state lifetimes at the indicated concentrations of eIF4B. Error bars represent the derived 95% CI of the mean lifetimes. From left to right: n = 292, 520, and 209.

Given the exceptionally slow association of eIF4A, we hypothesized that inclusion of other eIF4 proteins would accelerate eIF4A binding near the cap. eIF4A bound ∼10-fold more rapidly (22-42 s) in the presence of eIFs 4E, 4G, and 4B (Fig. 1d). Rapid eIF4A association depended on the concentration of eIF4F (Fig. S1c), consistent with a bimolecular interaction (0.50 ± 0.04 µM^−1^s^−1^) (Fig. 1c). We therefore conclude that these rapid eIF4A binding events reflect eIF4A binding within the eIF4F complex at or near the 5′ cap. In this context, the bound state lifetime was 14.2 ± 0.2 s (Fig. S1f), similar to that of eIF4A alone, and independent of eIF4F concentration (Fig. S1d).

Next, we determined how the composition of eIF4F modulated its binding. Omission of either eIF4G or eIF4E individually slowed the mean association times by ∼3-fold relative to the intact complex (Fig. 1d and Fig. S1e), with the bound state lifetimes unaffected (Fig. S1f). Thus, eIF4G and eIF4E work collaboratively to facilitate rapid cap-proximal association of the eIF4F complex, and once bound, eIF4A dwells on the mRNA independently of these cofactors. By contrast, omission of eIF4B had no effect on the association time (Fig. 1d and Fig. S1e), but the eIF4A bound state lifetime lengthened in an eIF4B concentration-dependent manner, extending modestly to 22.6 ± 0.4 s (Figs. 1e and S1h) at the concentration near the estimated micromolar-range equilibrium dissociation constant (*K*_D_) of eIF4A and eIF4B in yeast^29^. The eIF4A association time also doubled at the highest eIF4B concentration (Fig. S1g), demonstrating some inhibition of eIF4F cap-proximal binding by eIF4B. Thus, eIF4G and eIF4E rapidly target eIF4F to the 5′ cap, while eIF4B modulates eIF4A dissociation.

To determine the effects of ATP hydrolysis by eIF4A, we replaced ATP in our assay with saturating levels of either a non-hydrolyzable ATP analog (ADPNP), a slowly hydrolyzable analog (ATPγS), adenosine diphosphate (ADP), or removed the nucleotide entirely (apo-4A). Each condition strongly reduced the fraction of mRNAs showing eIF4A binding events at the cap (Fig. S1i), with the ADP and apo-4A conditions essentially eliminating observable FRET events. Consistent with their dramatic reduction in binding events, inclusion of ADPNP or ATPγS slowed eIF4F association by 3-fold (Fig. 1d and Fig. S1k) and reduced the eIF4A bound state lifetime by 1.5- to 2- fold (Fig. S1j). Considering that ATP hydrolysis is not required for eIF4F complex formation, these results demonstrate the importance of ATP hydrolysis by eIF4A for fast and stable eIF4F cap-proximal binding.

We observed transitions between states with different FRET efficiencies in a subset of complexes (Fig. S2a). Considering eIF4G contains two eIF4A binding sites, we hypothesized that multiple copies of eIF4A may bind proximal to the 5′ cap. To test this idea, we mixed equimolar amounts of eIF4A labeled with distinct FRET acceptor dyes (Cy5 and Cy5.5) in our single-molecule assay (Fig. S2b). If multiple copies of eIF4A simultaneously bound near the cap, this reaction scheme would deconvolute the multiple FRET states observed with Cy5-eIF4A alone into distinct binding events (Fig. S2c). As predicted, we detected two eIF4A molecules bound simultaneously proximal to the cap, as indicated by overlapping Cy5-eIF4A and Cy5.5-eIF4A binding events on individual mRNAs (Fig. S2c), with a second eIF4A association efficiency of 42% (Fig. S2f). Association of a second eIF4A molecule was 4-fold faster than the first (Fig. S2d), while the lifetime of the doubly bound state was comparable to that of the singly bound state (Fig. S2e). Moreover, removing any of the eIF4F/eIF4B cofactors greatly reduced the occurrence of double eIF4A binding events (Fig. S2f). Collectively, these results indicate that the eIF4F complex can stimulate the recruitment of multiple eIF4A molecules to the 5′ cap.

### Features of mRNA modulate eIF4F binding kinetics

To test the role of the cap in eIF4F binding to the mRNA 5′ region, we prepared and tested a β-globin mRNA lacking the cap structure but carrying a Cy3 dye at the 5′ end. As an additional control, we examined Cy3-cap labeled β-globin mRNA in the presence of saturating levels of 7-methylguanosine 5′-triphosphate (m^7^GTP) to compete with the eIF4E-cap interaction (Fig. 2a). Using our 5′ end-to-eIF4A FRET assay, eIF4F association to uncapped mRNA was 2-fold slower compared to capped mRNA (Fig. 2b). The addition of m^7^GTP to capped mRNA slowed eIF4F association 3-fold (Fig. 2b), comparable to the effect of omitting eIF4E. In both conditions, the eIF4A bound state lifetime remained unchanged from the capped mRNA control (Fig. 2c). Thus, the 5′ cap is a central element for rapid 5′ end targeting of the eIF4F complex, but not for subsequent stability of the eIF4A bound state.

**Figure 2.**
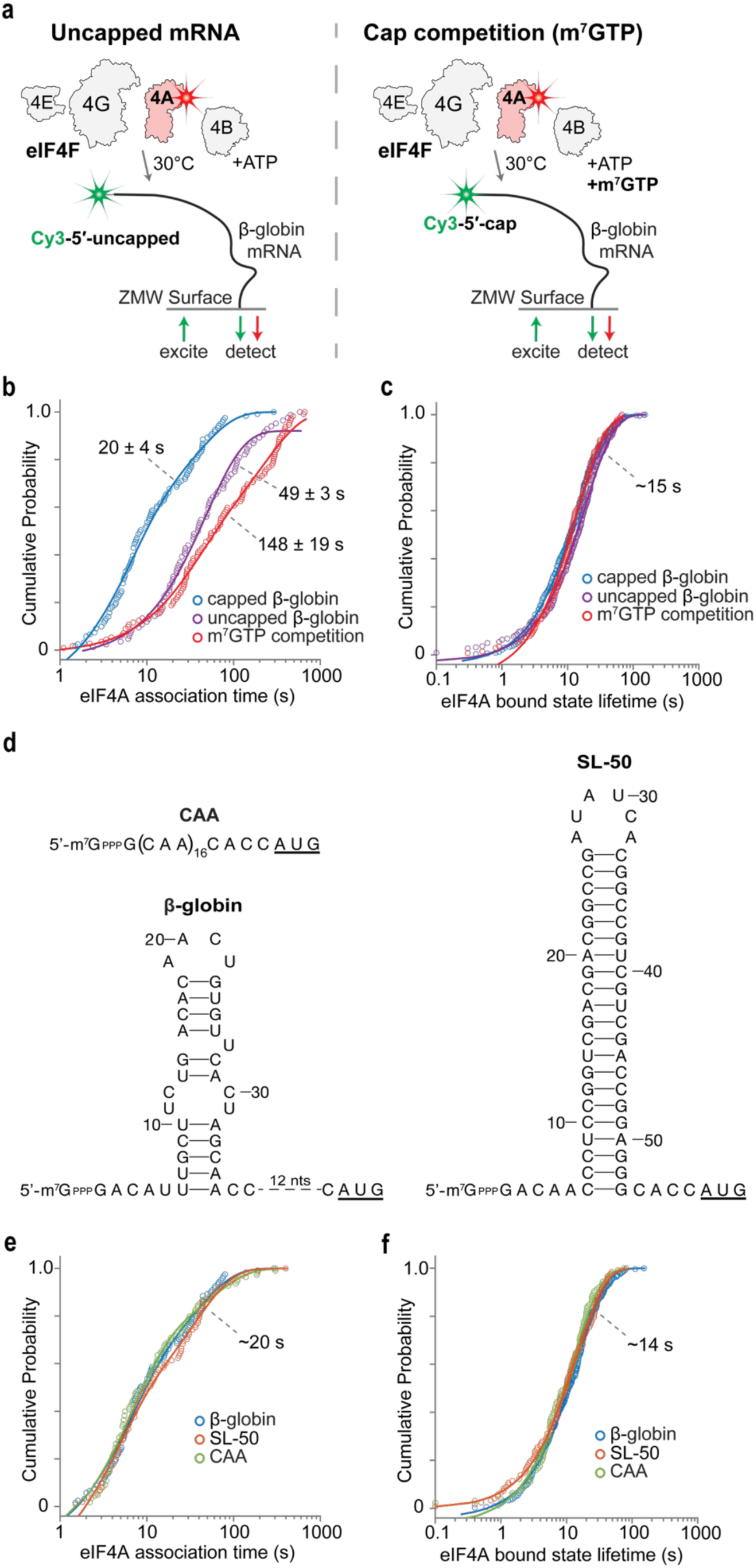
Features of mRNA modulate eIF4F binding kinetics. **a.** Schematics of single-molecule FRET assay for measuring Cy5-eIF4A binding to uncapped β-globin mRNA (left) or measuring binding in competition with m^7^GTP (right). **b, c.** Cumulative probability plots of the indicated parameters, comparing capped β-globin to the conditions denoted in (**a**). The lines represent fits to exponential functions, and the denoted values are derived mean times and 95% CI from the fit. For capped, uncapped, and m^7^GTP competition: n = 121, 141, and 142 respectively. **d.** Schematic of predicted RNA secondary structures in the 5′ UTR of the indicated mRNAs. **e.** Cumulative probability plots of the indicated parameters, comparing the different mRNA denoted in (**d**). The lines represent fits to exponential functions, and the denoted values are approximate mean times, which were comparable between all conditions. For SL-50 and CAA: n = 101 and 110 respectively.

Resolving mRNA secondary structure is critical for proper loading of the 40S ribosomal subunit, and highly stable secondary structures proximal to the 5′ cap inhibit translation^57,58^. RNA structures could disrupt eIF4F binding or downstream steps in translation initiation. To distinguish between these possibilities, we designed and labeled two capped model mRNAs containing either an unstructured sequence of CAA repeats (CAA mRNA) or a highly structured stem-loop with a predicted thermodynamic stability of -50 kcal/mol (SL-50 mRNA) upstream of the β-globin coding sequence (Fig. 2d). The stem-loop in SL-50 mRNA is situated 5 nt away from the 5′ cap, analogous to the predicted lower stability stem-loop (-10 kcal/mol) in the native β-globin mRNA 5′ UTR^59^ (Fig. 2d). In our single-molecule FRET assays, we observed identical eIF4F association times to the 5′ cap and eIF4A bound state lifetimes on the three distinct 5′ UTRs (Figs. 2e and 2f). These results indicated that rather than control eIF4F association, the cap-proximal structures (∼5 nt away) impact a step downstream of its initial eIF4F binding.

### Remodeling of mRNA by eIF4F and eIF4B is fast on the timescale of translation

To explore directly how eIF4F and eIF4B modulate mRNA 5′ end conformations, we developed a single-molecule FRET assay to monitor the relative proximity of the 5′ cap to internal positions in the mRNA (Fig. S3a). A folded RNA structure should be more compact, bringing the cap closer to downstream RNA segments. After testing multiple probe positions (Fig. S3b), we observed that annealing a Cy5-labeled DNA oligonucleotide probe within the coding sequence (124 nt from the cap) predominantly yielded high FRET efficiency with Cy3-cap-labeled β-globin mRNA (Figs. 3b and S3c). The majority of the mRNAs yielded FRET (58%), and the FRET state was highly stable (199 ± 9 s) and limited by the photostability of the dyes (Figs. S3d, e). The observed stable compact folded state of β-globin mRNA is consistent with predicted secondary structure in the 5′ UTR^59^.

**Figure 3.**
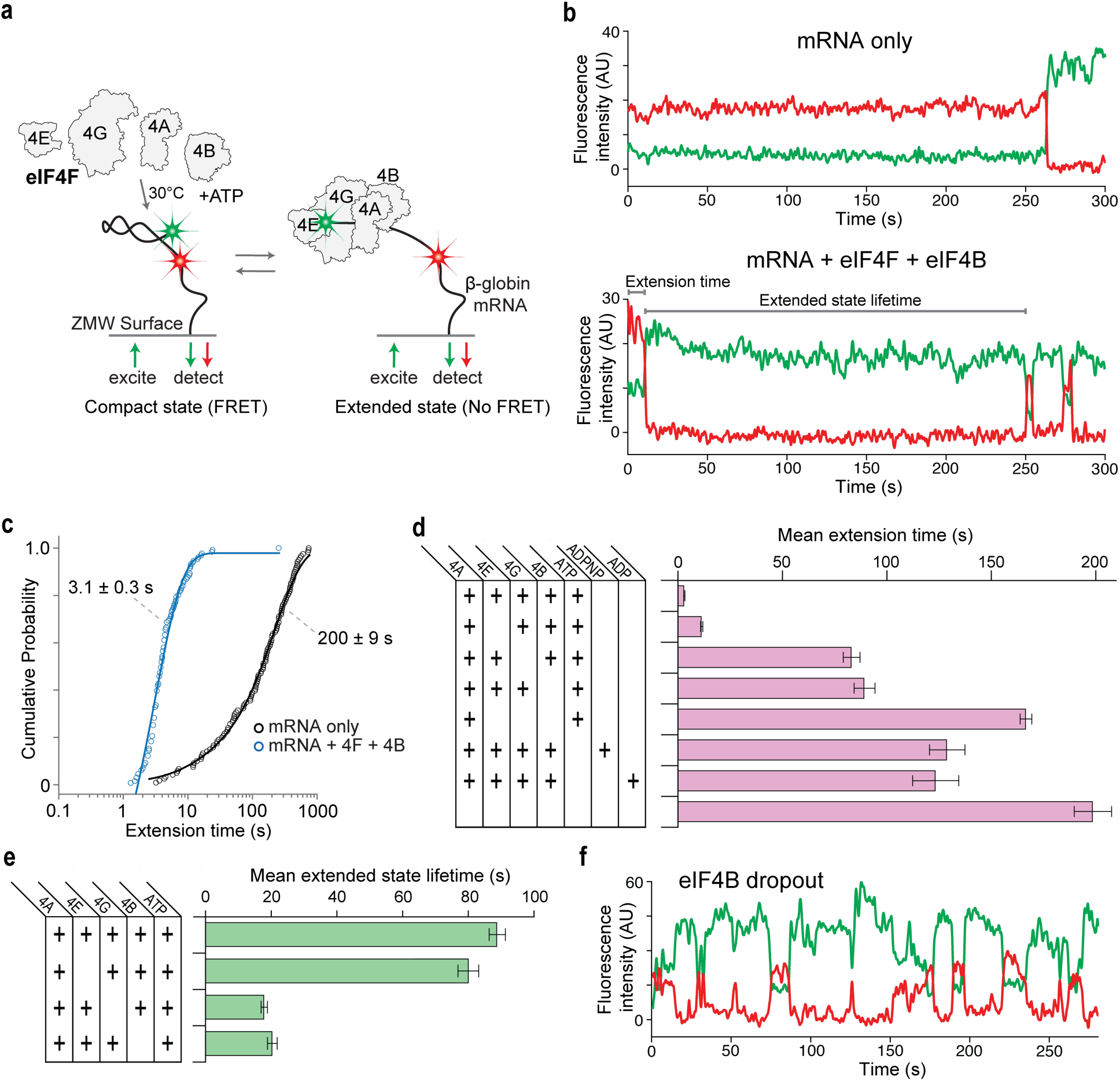
Remodeling of mRNA by eIF4F/eIF4B is fast on the timescale of translation. **a.** Schematic of single-molecule cap-to-probe FRET assay where eIF4A, eIF4E, eIF4G, eIF4B, and ATP are delivered to a Cy3-5′-cap-labeled β-globin mRNA with Cy5-probe 124 nt from the cap. Schematic denotes the two observed FRET states. **b.** Example single-molecule data for the assay. The top plot demonstrates the characteristic FRET in the absence of proteins. The bottom demonstrates the transition between FRET (compact) to loss of FRET (extended) in the presence of eIF4F/eIF4B. The lifetime of the first FRET state was denoted as extension time. Time to subsequent FRET was denoted as the extended state lifetime. **c.** Cumulative probability plot of the extension time, comparing the conditions denoted in (**b**). The lines represent fits to exponential functions, and the denoted values are derived mean times and 95% CI from the fit. **d, e.** Plots of mean extension times and mean extended state lifetimes across various conditions. Error bars represent the derived 95% CI of the mean times. From top to bottom: n = 131, 94, 106, 103, 117, 124, 111, and 121. **f.** Example single-molecule data for the eIF4B dropout condition showcasing the dynamic interchange between extension and compaction.

We next examined how eIF4F and eIF4B remodel mRNA conformation using this cap-to-probe FRET assay (Fig. 3a). Real-time addition of unlabeled eIF4A and ATP did not perturb mRNA conformation (Fig. 3d and Fig. S4a). In stark contrast, addition of complete eIF4F, eIF4B, and ATP reduced the lifetime of the initial FRET state by nearly 100-fold (3.1 ± 0.3 s) (Figs. 3b and 3c). This effect was consistent across all probe positions tested (Fig. S3i). Importantly, we found that the Cy5-oligonucleotide probe remained annealed to the tethered mRNA, with any minor mRNA-probe duplex unwinding occurring at least 10-fold slower than mRNA extension (Fig. S3h). We therefore conclude that the rapid loss of cap-to-probe FRET arose due to remodeling of the mRNA by eIF4F into conformations incompatible with FRET. We defined this state as the extended state and the elapsed time for it to be achieved as the extension time. The extended state was long-lived, persisting for 89 ± 3 s before the mRNA compacted again to yield a subsequent instance of cap-to-probe FRET (Fig. S3f). These compaction events were brief, with re-extension occurring in 4.3 ± 3 s (Fig. S3g). Dilution of eIF4F and eIF4B by 10-fold lengthened the extension time by 30-fold, as expected since the extension time also encapsulates the time for eIF4F to bind the mRNA. eIF4F dilution also shortened the extended state lifetime by 3-fold and lengthened the re-extension time by 4-fold (Figs. S3j-l). Furthermore, substitution of ATP with either ADPNP or ADP lengthened the extension time by more than 40-fold (Fig. 3d and Fig. S4d). Thus, eIF4F rapidly drives the mRNA into a long-lived extended state, which requires the ATP hydrolysis activity of eIF4A and depends on eIF4F and eIF4B concentration. The rates and rate constants used to derive mean times are listed in Table S2.

To understand how the composition of eIF4F and eIF4B contributed to mRNA remodeling, we systematically omitted eIF4 components. Excluding eIF4G lengthened the extension time by ∼30-fold (Fig. 3d and Fig. S4a) and decreased the extended state lifetime by 5-fold (Fig. 3e and Fig. S4b), consistent with its role as the scaffold for eIF4F and regulator of eIF4A activity. Omitting eIF4B yielded nearly identical effects, in agreement with its role as an eIF4A activator (Figs. 3d, e and Figs. S4a, b). In both cases, FRET efficiencies became more heterogeneous, with the mRNA sampling additional conformational states (Fig. 3f), suggesting inefficient and less coordinated remodeling by eIF4A. In contrast, omission of eIF4E only modestly lengthened the extension time by 4-fold and did not affect the extended state lifetime (Figs. 3d, e and Figs. S4a, b). Re-extension after subsequent compaction remained unchanged throughout all omissions (Fig. S4c). Collectively, our findings indicate that both eIF4G and eIF4B markedly enhance ATP-dependent remodeling of mRNA conformation by eIF4A, with eIF4E likely enhancing cap-proximal remodeling by promoting cap binding of eIF4F.

### Features of an mRNA can modulate its conformational remodeling

To test how the 5′ cap affected mRNA remodeling, we repeated the extension assays above using Cy3-labeled uncapped β-globin mRNA (Fig. 4a). This mRNA yielded long-lived FRET with the oligonucleotide probe annealed within its coding sequence (124 nt from the 5′ end) (Figs. S5a, b); thus, the 5′ region of the mRNA adopts a stable, compact conformation independent of the 5′ cap. However, compared to the capped mRNA, addition of unlabeled eIF4F, eIF4B, and ATP to the uncapped mRNA lengthened the extension time by 7-fold and modestly reduced the extended state lifetime by about 2.5-fold (Figs. 4b, c). Re-extension times remained similar to those of capped mRNA (Fig. S5c). As a separate control, we included saturating levels of m^7^GTP competitor with the capped mRNA (Fig. 4a). This condition was analogous to omission of eIF4E (within 2-fold) for all parameters measured (extension time, extended state lifetime, and re-extension time) (Figs. 4b, c and S5c). Thus, the 5′ cap both accelerates eIF4F association and promotes eIF4F/eIF4B-driven remodeling of mRNA conformation.

**Figure 4.**
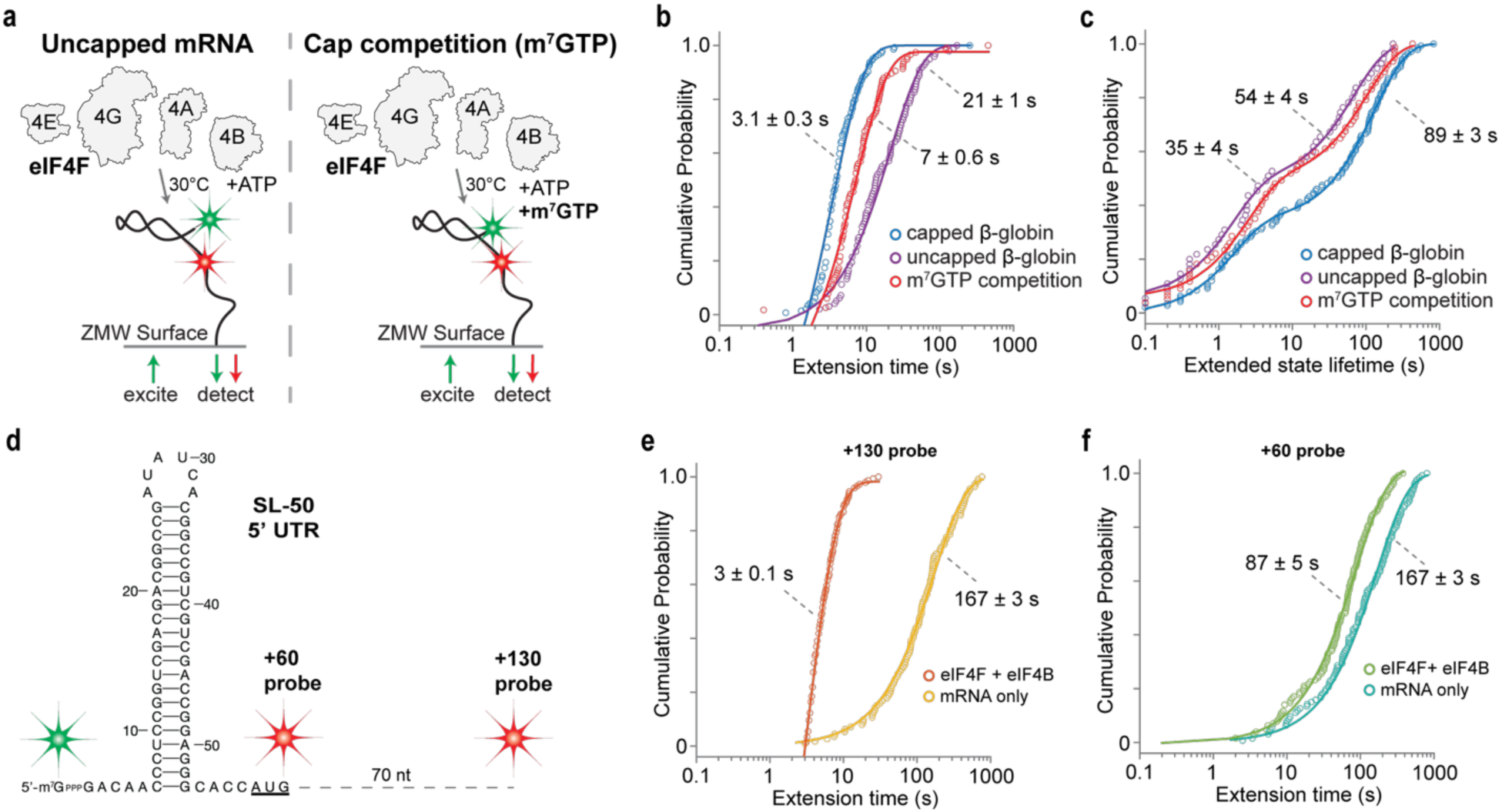
Features of an mRNA can modulate its conformational remodeling. **a.** Schematics of single-molecule FRET assay for measuring mRNA extension to Cy3-labeled uncapped β-globin mRNA (left) or measuring extension by eIF4F in competition with m^7^GTP (right). Cy5-probe is at position +124. **b, c.** Cumulative probability plots of the indicated parameters, comparing capped β-globin to the conditions denoted in (**a**). The lines represent fits to exponential functions, and the denoted values are derived mean times and 95% CI from the fit. For uncapped β-globin and m^7^GTP competition n = 98 and 116 respectively. **d.** Schematic of predicted RNA secondary structures in the 5′ UTR of SL-50 mRNA and the position of the different fluorescent probes utilized. The green star reflects the probe on the 5′ cap, the red stars denote the position of the Cy5 dyes on the different probes used. **e, f.** Cumulative probability plots of extension times comparing the effects of eIF4 proteins on extension of SL-50 mRNA, using the two probe positions shown in (**d**). The lines represent fits to exponential functions, and the denoted values are derived mean times and 95% CI from the fit. For mRNA only and eIF4F + eIF4B: in (**e**) n = 112 and 146, in (**f**) n = 126 and 141 respectively.

Since cap-proximal RNA structures inhibit translation initiation^57^ but had no effect on eIF4F binding (Figs. 2e, f), we next tested whether secondary structures inhibit eIF4F/eIF4B-mediated remodeling. We leveraged our Cy3-cap model mRNAs (CAA and SL-50) (Fig. 2d) that span the spectrum of secondary structure in our cap-to-probe FRET assay. We utilized the same Cy5-oligonucleotide probe directly matching our above β-globin experiments — located 124 nt from the cap in β-globin and CAA, and 130 nt from the cap in SL-50 on account of its slightly longer 5′ UTR (Fig. S5d). With this probe, the unstructured CAA mRNA rarely inhabited a stable FRET state, whereas SL-50 yielded a long-lived high-FRET state indicating a stable, compact conformation (Figs. S5e, f). Upon the addition of eIF4F, eIF4B, and ATP, SL-50 mRNA conformation extended rapidly, closely mirroring the rapid remodeling kinetics observed for β-globin (Fig. 4e). Given the distance from the annealed probe to the cap, this FRET signal may convolute both cap-adjacent and more distal RNA rearrangements. We therefore moved the probe to the 3′ base of the stable hairpin encoded in SL-50 mRNA (60 nt from the cap in SL-50), placing both Cy3-cap and Cy5-probe dyes within 6 nt of the base of the hairpin stem-loop (Fig. 4d). Using this hairpin-proximal signal, we observed a markedly lengthened extension time for the SL-50 mRNA, about 30-fold longer relative to the distal probe (Fig. 4f). In contrast, measured extension times using different probe positions were similar using β-globin mRNA (Fig. S3i). Collectively, our findings indicate that highly stable local secondary structures resist extension by eIF4F and eIF4B, despite similar resolution of weaker, longer-range interactions.

### The extended mRNA state is necessary for 43S PIC loading

Having shown that eIF4F rapidly remodels compacted mRNA into an extended state, we hypothesized that mRNA extension allows loading of the 43S PIC. We modified an established^43^ single-molecule FRET assay to monitor the recruitment of fluorescently labeled 40S ribosomes. We assembled the 43S PIC containing 40S subunits conjugated to a Cy3 dye and delivered the 43S PIC (5 nM) with molar excess eIF4F, eIF4B, and ATP simultaneously to tethered unlabeled capped β-globin mRNA (Fig. 5a). Loading of the 43S PIC was indicated by a burst of Cy3 fluorescence, and we determined the 43S PIC loading time relative to the delivery of components (Fig. 5b). Ribosomal loading displayed multi-step binding kinetics (Fig. 5c). The observed times were best described by a sequential two-step model, with a fast step of 9 ± 3 s and a slow step of 56 ± 2 s. The rates used to derive mean times are listed in Table S3.

**Figure 5.**
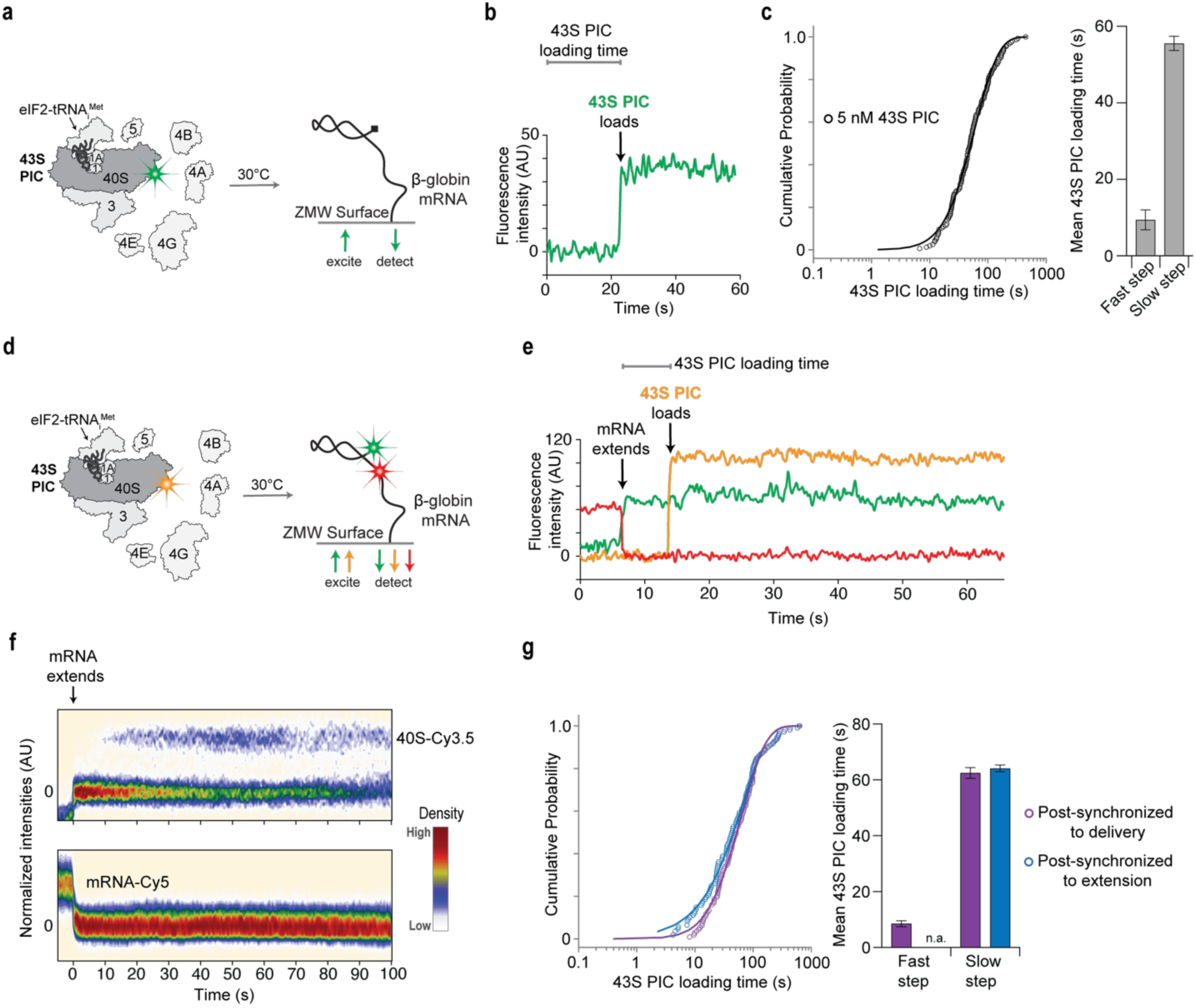
The extended mRNA state is necessary for 43S PIC loading. **a.** Schematic of single-molecule 43S PIC loading assay. All shown initiation factors and ribosomal subunits are delivered to a tethered β-globin mRNA in real-time. **b.** Example single-molecule data for the assay. The burst in green intensity represents 43S PIC loading and the time taken for this burst is the 43S PIC loading time. **c.** On the left, a cumulative probability plot of the experimental setup shown in (**a**). The line represents the fit to a hypoexponential function. On the right, a bar plot of the mean times for both kinetic steps derived from the fit. Error bars denote 95% CI from the fit, n = 156. **d.** Schematic of single-molecule mRNA extension and 43S PIC loading assay. Cy5-label on β-globin mRNA is at position +124 from the cap. **e.** Example single-molecule data for the assay in (**d**). The end of FRET between green and red denotes mRNA extension. The burst in yellow intensity represents Cy3.5-43S PIC loading. **f.** Heat maps of normalized intensities of 40S-Cy3.5 (top) and mRNA-Cy5 (bottom) for all analyzed events, n = 109. mRNA extension is labeled. **g.** On the left, cumulative probability plots of PIC loading post-synchronized to either delivery of factors (purple) or to mRNA extension (blue). The purple and blue lines represent the fits to exponential and hypoexponential functions respectively. On the right, a bar plot of the mean times for either two kinetic steps (purple) or single step (blue) derived from the fits. Error bars denote 95% CI from the fit.

To dissect what molecular events the two steps reflect, we examined 43S loading onto a capped mRNA preincubated with eIF4F and eIF4B (Fig. S6a), which allowed any mRNA activation steps to complete prior to 43S PIC addition. Under these conditions, a single-step model was sufficient to describe 43S PIC loading, yielding a mean loading time of 20 s (Fig. S6b). While this time did not match either the fast or slow step, the collapse of the multi-step binding kinetics into a single-step model suggested that at least one of the steps reflected mRNA selection and activation by eIF4F and eIF4B. Since eIF4A can bind directly to the 40S subunit, we hypothesized that the mixing of eIF4A and the 43S PIC prior to delivery could slow PIC loading. To test this, we performed the assay with eIF4F preincubated on the mRNA but with additional eIF4A pre-mixed with the 43S PIC (Fig. S6c) and found that PIC loading time in this condition indeed was 4.5-fold slower (Fig. S6d) and more closely aligned with the previously mentioned slow step. We thus hypothesized that the slow step correlates to PIC loading on an activated mRNA, whereas the fast step correlates to the mRNA activation steps. The similar magnitude to the mRNA extension times previously observed supports this hypothesis (Fig. 3c).

We hypothesized that combined eIF4F/4B binding and ATP-driven extension of β-globin mRNA was the fast step observed in our 43S PIC loading assay. If valid, 43S PIC loading times measured relative to mRNA extension should collapse into a single-step binding model. We simultaneously monitored mRNA extension by eIF4F and 43S PIC loading by combining our cap-to-probe FRET and 43S PIC loading single-molecule assays. We tethered to the imaging surface the Cy3-capped β-globin mRNA annealed with the Cy5-oligonucleotide probe and in real time we added eIF4F, eIF4B, ATP, and the 43S PIC labeled on the 40S subunit with Cy3.5 dye (Fig. 5d). As above, mRNA extension was indicated by loss of Cy3-cap to Cy5-probe FRET, and 43S PIC loading by a burst of Cy3.5 fluorescence (Fig. 5e). Relative to reagent delivery, we observed mean extension times and two-step 43S PIC loading times consistent with our findings in the individual assays (Fig. 5g and Figs. S6e, f). Strikingly, though, 43S PIC loading times measured relative to initial mRNA extension fit a single-step model with a mean time of 64 ± 1 s (Fig. 5g), matching the slow step in the two-step model. Furthermore, almost all 43S PIC loading events (99%) on β-globin mRNA occurred during its extended state (Fig. 5f and Fig. S6g). Omission of eIF4F decreased 43S PIC loading by over 90%, further consistent with an inability of the 43S PIC to load onto the compact state (Fig. S6g). Collectively, these results demonstrate that the fast step in PIC loading correlates to mRNA selection and extension by eIF4F and eIF4B, suggesting this as the activation step necessary for the 40S subunit to load, while the slow step represents 43S PIC loading to an activated mRNA, at our limiting concentration of 43S PIC.

### Regulation of 43S PIC loading by eIF4F composition and mRNA features

Our proposed model yielded several testable predictions. First, increasing 43S PIC concentration should accelerate only the slow step (40S loading time to an activated mRNA), whereas the fast step (mRNA activation time) should remain unchanged. In agreement, using our 43S PIC loading assay (Fig. 5a), we found that increasing 43S PIC concentration specifically hastened the slow step (bimolecular rate constant of 2.6 ± 0.7 µM^−1^s^−1^) with minimal effect on the fast step (Fig. 6a and Figs. S7a, S7j). Second, since mRNA remodeling by eIF4F/eIF4B was sensitive to cap-proximal secondary structures (Fig. 4f), the 5′ cap (Figs. 4b, c), and temperature (higher temperature accelerates extension) (Figs. S7b-d), these features should modulate the fast step to a greater degree than the slow step. Consistently, increasing temperature from 30 °C to 37 °C in our 43S PIC loading assay accelerated the fast step ∼2-fold without affecting the slow step (Fig. S7e). In contrast, the highly structured 5′ UTR of SL-50 mRNA lengthened the fast step 2.5-fold relative to β-globin mRNA and our unstructured model (CAA) (Fig. 6c and Fig. S7h). Elimination of the 5′ cap also lengthened the fast step by 5-fold (Fig. 6d and Fig. S7i). However, on all four mRNA templates we examined, the slow steps of the two-step 43S PIC binding model were identical. These findings indicate that while mRNA extension is delayed by stable secondary structure or the absence of a cap, the 43S PIC loads onto the extended mRNA at the same rate, regardless of how structured the mRNA was prior to extension.

**Figure 6.**
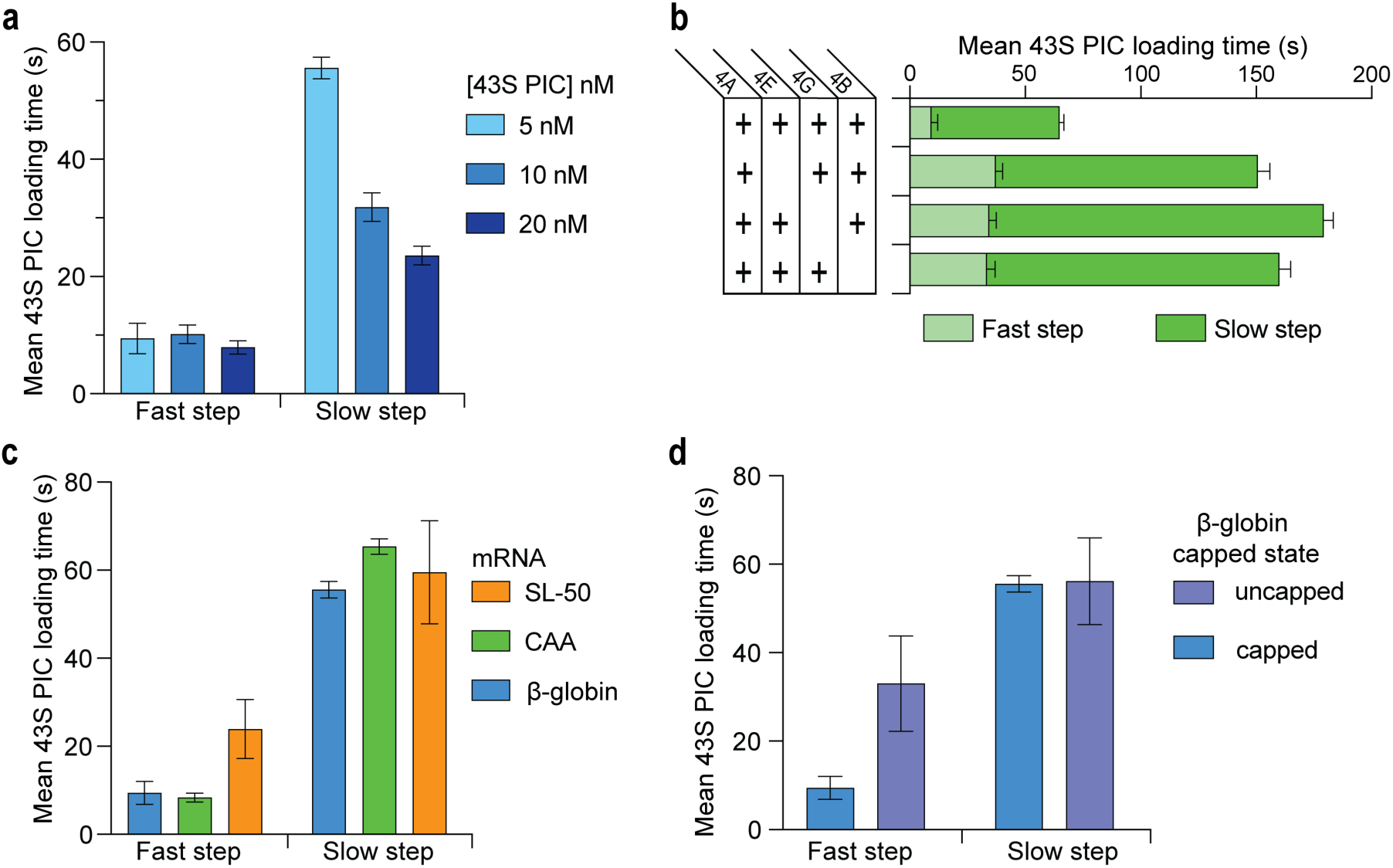
Regulation of 43S PIC loading by eIF4F composition and mRNA features. **a.** Bar plot of derived mean times for both kinetic steps (fast and slow) of 43S PIC loading at the indicated concentrations of 43S PIC. Error bars denote 95% CI from the fit, for 5, 10, 20 nM: n = 156, 119, and 98 respectively. **b.** Stacked bar plot of derived mean times of both kinetic steps (fast and slow) of 43S PIC loading for various conditions. Error bars denote 95% CI from the fit. From top to bottom n = 156, 118, 118, and 115. **c.** Bar plot of derived mean times for both kinetic steps (fast and slow) of 43S PIC loading with the indicated mRNAs. Error bars denote 95% CI from the fit, for CAA and SL-50: n = 160 and 120 respectively. **d.** Bar plot of derived mean times for both kinetic steps (fast and slow) of 43S PIC loading with the capped state of β-globin mRNAs. Error bars denote 95% CI from the fit, for uncapped mRNA: n = 105.

Lastly, we predicted that the concentration and composition of eIF4F and eIF4B should impact both the fast and slow steps of the two-step 43S binding model, as individual eIF4 components actively remodel the mRNA and recruit the 43S PIC. Indeed, reduction of the eIF4F concentration by 10-fold delayed the fast and slow steps by 2-3-fold (Fig. S7f). Similarly, omission of eIF4G, eIF4E, or eIF4B slowed the fast step by ∼4-fold and delayed the slow step 2-fold (Fig. 6b and Fig. S7g). Together, these findings agree with the delayed and disrupted mRNA extension times observed above (Figs. 3d, e) and strongly support a role for each component in 43S PIC loading.

## Discussion

Despite decades of biochemical and genetic studies that developed our understanding of the eIF4F complex and its role in translation initiation^1,2,4,7^, the nature of mRNA activation by these proteins has remained vague. Here, we defined a mechanistic framework for the early steps in translation initiation by characterizing the binding of eIF4F to the 5′ end of an mRNA, the subsequent remodeling of mRNA conformation, and how this remodeling promotes the loading of the 43S PIC.

Efficient loading of the 40S ribosomal subunit to the 5′ end requires mRNA activation by the eIF4F complex. Consistent with established models in which eIF4E and the eIF4G scaffold target the complex to the 5′ cap^7,12,13,17^, we found that both cofactors and the 5′ cap collaboratively accelerate eIF4A association to the cap-proximal region. After initial eIF4A binding, a second eIF4A can associate, which we posit involves the second eIF4A-binding domain within eIF4G^60^. Whereas eIF4E and eIF4G primarily enhanced eIF4A association, eIF4B uniquely increased eIF4A residence time on the mRNA. Prior studies demonstrate that cap binding stabilizes the eIF4G:E subcomplex on an mRNA at timescales much longer than we observed for eIF4A binding^19^, suggesting that cap-bound eIF4G:E persists through multiple rounds of eIF4A binding and dissociation. In the presence of all eIF4 proteins and ATP, our single-molecule data yield an apparent cap-proximal eIF4A-mRNA *K*_D_ of ∼140 nM, consistent with previous bulk biochemical measurements^28^.

The β-globin mRNA 5′ region exhibits a stable compact conformation. Previous work on probing end-to-end interactions in β-globin mRNA demonstrated globally compact conformations^61^. However, end-to-end interactions were transiently formed for just seconds, while in this study we observed intermediate-range interactions stable for minutes, consistent with stable base pairing. Yet, upon binding to the 5′ end, eIF4F and eIF4B rapidly remodel the observed compact state of β-globin mRNA into an extended state. Although our mRNA extension and eIF4F association times cannot be directly compared – unlabeled eIF4A copurifies with eIF4G, slowing the observed association times – extension times show cap dependency, suggesting that mRNA remodeling is downstream of eIF4F cap-proximal binding. Given the rapid extension observed, we therefore hypothesize that the rate limiting step in β-globin mRNA remodeling is eIF4F binding, with mRNA extension, post binding, likely occurring rapidly (timescale < 1s).

Rapid RNA extension and stabilization of the extended state was strongly dependent on eIF4G, eIF4B, and ATP with relatively weaker contributions by eIF4E and the 5′ cap. Our results are consistent with prior work demonstrating eIF4A alone is a weak and non-processive RNA helicase^36^, and that eIF4G and eIF4B enhance eIF4A ATPase and RNA helicase activity^23,24,26,27^. Both also contain RNA recognition motifs that could potentially help stabilize the extended state of an mRNA^16,31^. The contribution of eIF4E and the 5′ cap to mRNA extension is likely by targeting the eIF4F complex to the 5′ end^28^ and position the complex in a conformation that is conducive to efficient mRNA remodeling, consistent with cap-mediated stabilization of eIF4F previously observed^19^. Additionally, eIF4E enhances the eIF4G-driven activation of eIF4A^25^, which could help stabilize the extended mRNA. We hypothesize that the multiple eIF4A binding events could produce a dynamic exchange whereby at least one eIF4A molecule remains bound constantly sustaining the extended state, likely facilitated by eIF4G and eIF4B. This is supported by the extended state lifetime being over 6-fold longer than eIF4A binding lifetimes. However, with eIF4G/eIF4B omission the extended state lifetime drops down to match a single eIF4A binding lifetime. The need for this dynamic exchange could also explain why eIF4A is the most abundant initiation factor^62,63^.

ATP was critical for both eIF4A-mRNA binding and subsequent mRNA extension. Replacing ATP with non-hydrolyzable analogs, ADP, or omitting ATP entirely largely destabilized both eIF4A association and binding lifetimes, consistent with its role in eIF4A-mRNA binding^24,28,35^. We note that in contrast to previous reports where these analogs delay departure of DEAD-box helicases from an mRNA^64^, we see faster departure, yet this is consistent with other measurements showing that these analogs reduce mRNA binding^35^. Finally, these ATP analogs also prevented extension consistent with duplex unwinding inhibition by ADPNP observed in similar DEAD-box helicases.^35,64^ Thus, ATP free energy is used by eIF4A in conjunction with the eIF4G and eIF4B to create and maintain the extended RNA state, and to allow subsequent dissociation of eIF4A for factor exchange.

Extension of mRNA conformation by eIF4F gates 43S PIC loading. In structures of mRNA-bound ribosomes, mRNA is single-stranded within the binding channel on the 40S subunit^34,40^. Secondary structure, and consequently, the compact conformation could thus pose a steric barrier to loading of the 40S subunit. We observed multistep binding behavior for 43S PIC loading, which fit a two-step kinetic model. By tracking mRNA extension alongside 43S PIC loading, we were able to deconvolute the two steps and established that the faster step was initial mRNA extension by eIF4F/eIF4B and the slower step was bimolecular 43S PIC loading to an extended mRNA. A 43S PIC struggles to load onto compact mRNA states, but an extended mRNA acts as an efficient bimolecular substrate for the PIC.

Aside from promoting efficient mRNA remodeling, eIFs 4G, 4E, and 4B also mediate 43S PIC loading after initial extension, consistent with eIF4G binding directly to eIF3^14^ or eIF4B and eIF4G binding to the 40S subunit^38,41^ on the 43S PIC. While our assay tracks initial 43S loading, recent work outlines the subsequent mRNA accommodation mechanism^65^. In that framework, accommodation rates resemble our slower loading step when using higher 43S PIC concentrations, suggesting that our limiting PIC conditions partition the activation step from subsequent loading. Local eIF4F, 43S PIC, and mRNA availability may therefore determine whether mRNA activation and loading are temporally decoupled or proceed as a kinetically coupled event.

Our results are consistent with a simple kinetic model (Fig. 7) of eIF4F-mediated 43S PIC loading that begins with eIF4F binding to the 5′ cap. The components within eIF4F facilitate fast recruitment to the 5′ end through interactions with the 5′ cap, while eIF4B stabilizes eIF4A binding. The association of eIF4F likely functions as the rate-limiting step in mRNA remodeling, as once bound, mRNA conformation is rapidly extended. Extension is strictly dependent on eIF4A-catalyzed ATP hydrolysis and strongly promoted by the other eIF4 proteins, which help maintain mRNA in an extended state, and likely remain bound to promote fast 43S PIC recruitment. This activated state serves as a necessary precursor to 43S PIC loading by providing an extended mRNA amenable to docking into the mRNA binding channel on the 40S subunit and an anchor in eIF4G to drive loading through interactions with eIF3. The requirement for extension prior to 43S PIC loading could hint towards a slotting model^5^ of 43S PIC loading, where a multi-nucleotide stretch of unwound mRNA is necessary to insert properly and laterally into the 40S subunit binding channel.

**Figure 7.**
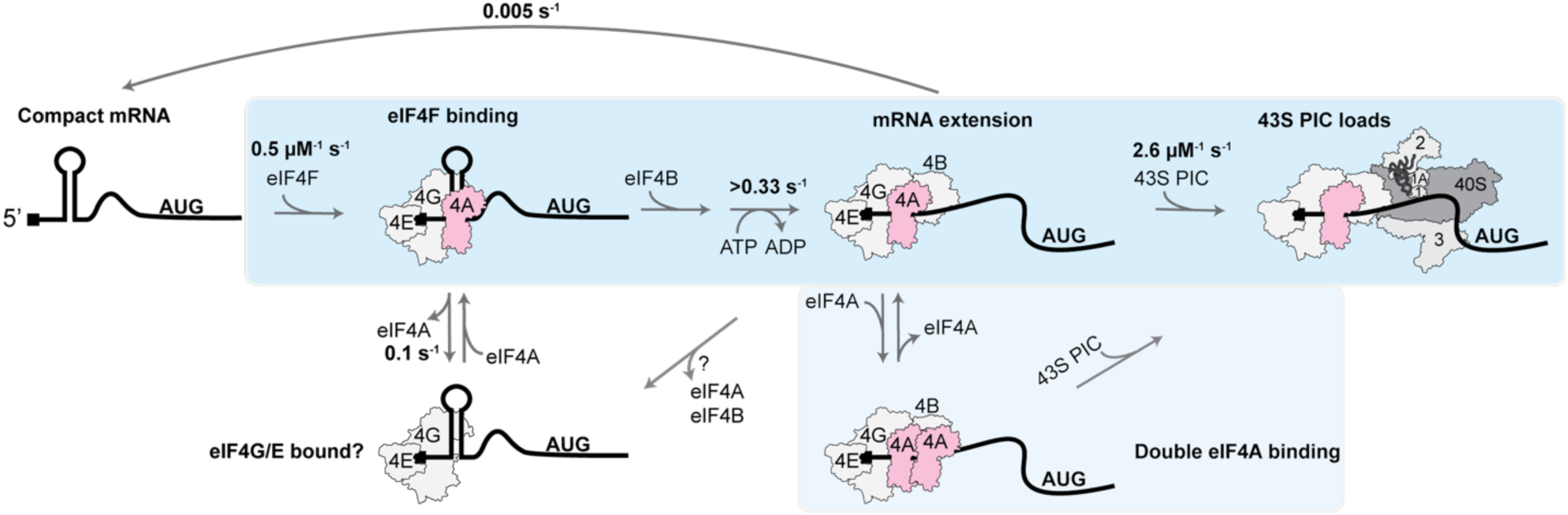
Kinetic model for the early steps of translation initiation. On the left, compact mRNA is represented as a cartoon with a 5′ cap, secondary structure, and start codon. Blue backgrounds denote on-pathway steps. In the first step, eIF4F binds to the 5′ end of an mRNA, guided by the 5′ cap and facilitated by eIF4G/E. eIF4A dissociates independently of eIF4F, with eIF4G/E possibly remaining bound. Upon initial binding of eIF4F, mRNA conformation is very rapidly remodeled into an extended conformation in an ATP-dependent manner, with eIF4B likely binding prior to this rapid extension step. This extended state is stabilized by the proteins, with re-compaction occurring slowly. Critically, the extended state is the amenable conformation for 43S PIC loading at the 5′ end. We speculate that the extended state is maintained by the continuous presence of at least one eIF4A molecule at the 5′ end. This continuity is likely sustained by the overlapping binding cycles of multiple eIF4A molecules.

Our kinetic model provides timescales for the early steps of initiation on mRNAs in human cells. Using reported binding affinities of the eIF4G scaffold to eIFs 4E and 4A^23,60,66^, we estimate cellular concentrations of the eIF4F complex to be in the range of ∼300 nM^63^, about 3-fold higher than those used in our mRNA extension assay. At such concentrations, we estimate eIF4F cap-binding and subsequent extension to occur within 1 s, whereafter the 43S PIC can load. At cellular concentrations of the 43S PIC (∼1.5 µM, estimated from concentration of limiting reagent – eIF3^63^), our 43S PIC binding rate constant (2.6 µM^−1^ s^−1^) indicates loading takes ∼0.3 s. Collectively, we show the early steps of initiation would occur in under 2 s, well within the ∼30 s time scale of full initiation previously reported, with eIF4F cap-proximal binding as the rate limiting step in ribosomal loading. Rapid recognition, extension, and 43S PIC loading on mRNAs with less stable secondary structures in their 5′UTR might explain their high translation efficiencies^67–69^. In contrast, mRNA with highly stable cap-proximal secondary structures had similar eIF4F binding kinetics but much slower mRNA extension that delayed 43S PIC loading, consistent with previous studies demonstrating cap-proximal structures inhibit translation^57^. In such mRNAs, the rate-limiting step appears to be conformational remodeling after factor binding. Likewise, changes in eIF4F/eIF4B and 43S PIC availability through cell signaling or in disease states would also affect these rates, therefore modulating the efficiency of translation^46,48^.

The results presented here establish a biophysical and mechanistic framework for the critical process of mRNA selection and activation in protein synthesis, yet questions remain. The nature of the extended state remains unclear. Our results show stable hairpins near the 5′ end, such as in SL-50, are likely not unwound, whereas weak secondary structures and longer-range mRNA conformation were rapidly extended by eIF4F/eIF4B. 43S PIC loading rates after the delayed mRNA extension proceeded at a comparable rate to unstructured mRNA, suggesting cap-proximal structure might not need to be completely unwound for subsequent 43S PIC loading but may require a shorter unwound stretch amenable to ribosomal docking. The coupling of these early steps of initiation to downstream scanning and start codon recognition must be investigated in future work.

## Methods

### Labeled eIF4A

A synthetic DNA encoding for full-length human eIF4AI with an N-terminal 6xHis tag, tobacco etch virus (TEV) protease cleavage site, and ybbR tag was purchased from Integrated DNA Technologies (IDT). The geneblock was inserted into a pET-28a plasmid backbone using restriction enzyme sites NcoI and XbaI. The plasmid was transformed into BL21 cells (Invitrogen #C600003) and grown overnight at 37 °C on Luria-Bertani (LB) agar plates containing 50 µg ml^−1^ kanamycin. Single colonies from the plate were grown at 37 °C into liquid cultures in LB broth containing 50 µg ml^−1^ kanamycin up to an optical density (OD) at 600 nM of 0.5. The liquid culture was then induced by 1 mM IPTG and grown for an additional 4 hours before cells were harvested by centrifugation at 5,000g for 20 min at 4 °C in a Fiberlite F9 rotor (Thermo Fisher, 13456093). Cells pellets were resuspended into lysis buffer (20 mM Tris-HCl pH 7.5, 300 mM KCl, 10% (v/v) glycerol, 10 mM imidazole, 5 mM β-mercaptoethanol) and lysed by sonication. The lysate was clarified through centrifugation at 38,000g for 30 min at 4 °C in a Fiberlite F21 rotor. Clarified lysate was loaded onto a Ni-NTA gravity-flow column that was equilibrated in lysis buffer. Upon loading, the column resin was washed with 10 column volumes (CV) of lysis buffer, 10 CV of wash buffer (20 mM Tris-HCl pH 7.5, 800 mM KCl, 10% (v/v) glycerol, 25 mM imidazole, 5 mM β-mercaptoethanol), and 10 CV of lysis buffer again. Protein was eluted from the column with elution buffer (20 mM Tris-HCl pH 7.5, 100 mM KCl, 10% (v/v) glycerol, 300 mM imidazole, 5 mM β-mercaptoethanol) into three fractions at 2.5 CV each. Fractions were screened through SDS-PAGE and relevant fractions were combined. Combined fractions were dialyzed overnight at 4 °C into TEV buffer (20 mM Tris-HCl pH 7.5, 250 mM KCl, 10% (v/v) glycerol, 5 mM β-mercaptoethanol) and simultaneously incubated in the dialysis cassette with TEV protease. Cleaved protein was separated from TEV protease through subtractive Ni-NTA gravity-flow column and clarified by syringe filtering using 0.2 µm filters. Protein was further purified through size-exclusion chromatography, loaded onto a Superdex 200 prep grade (pg) resin in HiLoad column, 26/600 (Cytiva #28989336) and eluted into fractions in ybbR labeling buffer (20 mM HEPES-KOH pH 7.5, 250 mM NaCl, 10 mM MgCl_2_, 10% (v/v) glycerol, 1 mM DTT). Fractions were screened through SDS-PAGE and fractions with ybbR-eIF4A purity >95% were combined. Purified ybbR-eIF4A was incubated at 37 °C for 4 hours with 4 µM Sfp synthase and 3x molar excess of Cy5-CoA or Cy5.5-CoA in the dark. Excess free dye was removed using a 10DG desalting column (Biorad #7322010) and the protein was further cleaned through size exclusion chromatography as described above and eluted into storage buffer (20 mM HEPES-KOH pH 7.3, 250 mM KOAc, 10 mM MgCl_2_, 10% (v/v) glycerol, 1 mM DTT). Labeling efficiency of eIF4A was around 50-75%, and labeled protein was frozen using liquid nitrogen and stored at -80 °C until use.

### Unlabeled eIFs

Recombinant eIF4AI^27,70^, eIF4E^25,27^, eIF4GI_165-1599_ ^43^, eIF4B^43^, eIF1^70^, eIF1A^70^, eIF3j^70^, and eIF5^71^ were purified as previously described. Endogenous human eIF2 and eIF3 were purified as previously described^70,72^. Human Met-tRNA_i_^Met^ was prepared as previously described^43^.

### 40S ribosomal subunits

Human 40S ribosomal subunits were purified from edited HEK 293T cell lines (RPS15-ybbR) and fluorescently labeled with either Cy3 or Cy3.5 as previously described^43^.

### Fluorescently labeled capped mRNAs

Linear DNA template for full-length β-globin mRNA (NM_000518.5, 628 nt) was prepared as previously described^43^. Plasmids encoding for CAA and SL-50 mRNAs on a PUC19 backbone were designed following similar templates previously described^57^ and ordered from GenScript. Linear DNA templates for CAA and SL-50 were obtained by linearizing the plasmids with restriction enzyme XbaI. All DNA templates used are listed in Table S5. All mRNA were in vitro transcribed and co-transcriptionally capped with N_3_-m^7^GpppG using T7 polymerase for 4 hours at 37 °C. The cap analog N_3_-m^7^GpppG was synthesized as previously described^56^. Transcription reactions were incubated with Turbo DNase (ThermoFisher # AM2238) for 15 minutes at 37 °C and transcribed mRNA were purified using MEGAclear Transcription Clean-Up Kit (ThermoFisher #AM1908). Purified mRNA was incubated with 1 mM Cy3-DBCO (Lumiprobe #213F0) in DBCO labeling buffer (30 mM HEPES-KOH pH 7.5, 1 mM EDTA, 2 mM Mg(OAc)_2_) for 40 minutes at room temperature in the dark. Excess free dye was removed using the transcription cleanup kit as described above. The Cy3-labeled mRNA was biotinylated at the 3ʹ end through potassium periodate oxidation and reaction with biotin-hydrazide as previously described^73^. The 5′ Cy3-labeled and 3ʹ biotinylated mRNA were further purified using the transcription cleanup kit as described above and stored at -80 °C until use.

### Fluorescently labeled uncapped mRNA

β-globin mRNA was *in vitro* transcribed with T7 polymerase and co-transcriptionally labeled at the 5′ end with guanosine-5′-(6-aminohexyl)-monophosphate (amino-GMP) (TriLink BioTechnologies #N-6009). The reaction was run for 4 hours at 37 °C after which TURBO DNase was added to digest the DNA template and transcribed mRNA were purified using MEGAclear Transcription Clean-Up Kit. 5′ amino-GMP labeled mRNA was incubated with Sulfo-Cy3-NHS (ThermoFisher # 24510) in HEPES pH 7.3 for 1 hour at room temperature in the dark. 5′ Cy3-labeled amino-GMP mRNA was biotinylated at the 3ʹ end as previously described^73^. The final 5′ Cy3-labeled, 3ʹ biotinylated mRNA was further purified using the transcription cleanup kit described above and stored at -80 °C until use.

### Non-fluorescently labeled mRNA

Linear DNA templates for all mRNA used were obtained as described above. β-globin mRNA was in vitro transcribed biotinylated at the 3ʹ end and enzymatically capped as previously described^43,73^. CAA and SL-50 mRNA were prepared just as β-globin mRNA up to the capping step, where both mRNA were enzymatically capped with the Faustovirus Capping Enzyme (NEB #M2081S) and 2’-O-methyltransferase (NEB #M0366) following the one-pot synthesis reaction protocol. All mRNA were purified using the MEGAclear Transcription Clean-Up Kit.

### Malachite green ATPase assay

The ATPase activity of eIF4A was tested via the liberation of free phosphates from ATP in the presence of mRNA using the Malachite green ATPase assay^74^. In assay buffer (20 mM HEPES-KOH, pH 7.3, 70 mM KOAc, 2.5 mM Mg(OAc)_2_), mixtures containing 1 µM of either eIF4A or ybbR-eIF4A were prepared. Separate mixtures were supplemented with 1 µM of eIFs 4E, 4G, and 4B to both eIF4A variants, and a control with no eIF4 proteins was included as well. These mixtures were incubated at 37 °C for 20 minutes after which 1 mM of ATP and 5 µg of yeast RNA (Roche #10109223001) were added and the reaction mixture was further incubated for another 30 minutes. A malachite green solution was added to the reaction mixture at 1:2 initial mixture to malachite green solution, giving the final concentrations of 200 µM malachite green, 2.3 mM ammonium heptamolybdate, 2.6% (v/v) concentrated H2SO_4_, and 0.03% (v/v) Tween-20. Reactions were incubated for 30 minutes at room temperature. Samples were tested for absorbance at 630 nm and full analyses were performed on Prism10 (Graphpad).

### Real-time single-molecule assays

All real-time single-molecule assays were performed using a modified Pacific Biosciences RSII DNA sequencer and the Maggie software (v. 2.3.0.3.154799) which was previously described^54^. Most experiments were performed at 30 °C except for the few indicated to be done at 37 °C. Excitation of Cy3 and Cy3.5 dyes was done using a 532-nm excitation laser at 0.32 µW µm^−2^, excitation of Cy5 and Cy5.5 dyes was done by either using a 642-nm laser at 0.1 µW µm^−2^ or through FRET as indicated. Movies were collected by detecting four-color fluorescent emission for all dyes (Cy3, Cy3.5, Cy5, and Cy5.5) at 10 frames per second for either 600 s or 900 s. Experiments were performed on zero-mode waveguide (ZMW) chips, purchased from Pacific Biosciences. Chips are washed with with 0.2% Tween-20 and TP50 buffer (50 mM Tris-acetate pH 7.5 and 100 mM KCl) prior to imaging. After washing, chips are incubated for 10 minutes with a neutravidin mixture containing 1 µM neutravidin, 0.7 mg ml^−1^ UltraPure BSA, and 1.3 µM of preannealed DNA blocking oligonucleotides (CGTTTACACGTGGGGTCCCAAGCACGCGGCTACTAGATCACGGCTCAGCT and AGCTGAGCCGTGATCTAGTAGCCGCGTGCTTGGGACCCCACGTGTAAACG) resuspended in TP50 buffer. After incubation, chips are washed thoroughly with TP50, leaving behind a neutravidin coated ZMW imaging surface.

A 4BGE mixture containing 11 µM eIF4B, 6.5 µM eIF4G, and 8 µM eIF4E is prepared in reconstitution buffer (20 mM HEPES–KOH pH 7.5, 70 mM potassium acetate, 2.5 mM magnesium acetate, 0.25 mM spermidine, 0.2 mg ml^−1^ creatine phosphokinase), when omission of these proteins is indicated, they are not included in this 4BGE mixture. To prepare the eIF2-GTP-Met-tRNA_i_ ternary complex (TC), 3.3 µM eIF2 was incubated in reconstitution buffer, supplemented with 1 mM GTP and additional creatine phosphokinase (final of 0.5 mg ml^−1^), for 10 min at 37 °C to saturate eIF2 with GTP. The eIF2-GTP complex was then incubated with 2.3 µM of Met-tRNA_i_ for 5 min at 37 °C to form the TC. To prepare the 43S PIC, 240 nM Cy3 or Cy3.5 (as indicated) labeled 40S subunits are incubated with 1 µM eIF1, 1 µM eIF1A, 500 nM ternary complex (by eIF2), 1 µM eIF5, 400 nM eIF3, 1.2 µM eIF3j, 1 mM ATP, and 1 mM GTP for 5 minutes at 37 °C in reconstitution buffer.

For all experiments, the indicated 3ʹ biotinylated mRNA was tethered to the neutravidin-coated ZMW surface. On mRNA with annealed fluorescent oligonucleotide probes, hybridization is performed prior to mRNA tethering by incubating the mRNA with molar excess of the indicated Cy5-labeled probe (all oligonucleotides purchased from IDT, and listed in Table S6) in 50 mM Tris-HCl and 100 mM KOAc for 2 minutes at 65 °C then snap cooled on ice. The mRNA or mRNA-probe complex were diluted in reconstitution buffer and incubated on the neutravidin-coated chip for 10 minutes. After mRNA immobilization, the chip is washed with reconstitution buffer to remove excess untethered mRNA.

After mRNA tethering, the chip surface is prepared with imaging buffer, reconstitution buffer supplemented with casein (62.5 µg ml^−1^), 5 mM TSY and an oxygen-scavenging system^75^ composed of 2 mM protocatechuic acid and 0.06 U per µl protocatechuate-3,4-dioxygenase. Unless otherwise specified, in experiments where eIF4A is included, the imaging buffer is supplemented with 1 mM ATP or an indicated ATP analog (ADPNP, ATPγS, or ADP). For experiments with cap competition, 1 mM m^7^GTP is supplemented to the imaging buffer. When the 43S PIC is included the imaging buffer is further supplemented with 1 mM guanosine triphosphate (GTP).

In the 5′ end-to-eIF4A FRET assay, a delivery mixture containing Cy5-labeled or Cy5.5-labeled eIF4A, or both is prepared in imaging buffer supplemented with the 4BGE mixture and incubated at room temperature for 10 minutes. Upon the start of the data acquisition, the delivery mixture is added in real-time to the mRNA tethered imaging surface resulting in a final concentration of 100 nM labeled eIF4A, 130 nM eIF4G, 160 nM eIF4E, and 220 nM eIF4B unless otherwise specified.

In the cap-to-probe FRET assay, the delivery mixtures were prepared as described above except unlabeled eIF4A replaced fluorescently labeled eIF4A. Upon the start of the data acquisition, the delivery mixture is added in real-time to the mRNA tethered imaging surface resulting in a final concentration of 1 µM unlabeled eIF4A, 130 nM eIF4G, 160 nM eIF4E, and 220 nM eIF4B unless otherwise specified. On conditions where no protein addition is indicated, the delivery mixture is prepared from only imaging buffer and added upon the start of data acquisition.

In the 43S PIC loading assays, the delivery mixture is prepared by adding eIF4A, the 4BGE mix, and 43S PIC to imaging buffer. Upon the start of the data acquisition, the delivery mixture is added in real-time to the mRNA tethered imaging surface resulting in a final concentration of 1 µM unlabeled eIF4A, 130 nM eIF4G, 160 nM eIF4E, and 220 nM eIF4B, 10 nM labeled 40S subunits, 20 nM ternary complex, 16 nM eIF3, 48 nM eIF3j, 290 nM eIF1A, 290 nM eIF1 and 290 nM eIF5 unless otherwise specified. In experiments indicated as eIF4F preincubated, all eIF4 proteins are omitted from the delivery mixture and instead supplemented to the imaging buffer on the ZMW chip surface and incubated with the tethered mRNA for 10 minutes at room temperature^43^. All final concentrations of initiation factors remain the same.

### Single-molecule data analysis

Experimental movies captured on the PacBio RS II instrument that contain fluorescent intensities over time were processed on MATLAB 2024a as previously described^37,43,54,73,76^. From the movies, fluorescent spots that correspond to ZMW wells were extracted by filtering for fluorescence intensity using custom MATLAB scripts. Extracted fluorescent traces plot fluorescence intensity of the indicated dye over time. For FRET data acquired from 5′ end-to-eIF4A and cap-to-probe FRET assays, fluorescent traces were processed on SPARTAN version 3.7.0^77^. On SPARTAN, traces were filtered by FRET lifetime greater than zero and Pearson’s correlation coefficient between donor and acceptor fluorescence between −1.1 and 0. Extracted traces were manually curated to select for events which contain single photobleaching events and clear FRET signals. Fluorescent background and donor-acceptor crosstalk were corrected on SPARTAN and by custom MATLAB scripts. Corrected traces were reformatted to be compatible with vbFRET version June10^78^, where vbFRET was utilized to automatically assign FRET states. Assigned FRET states were manually curated using custom MATLAB scripts, where states were selected as either FRET-on or FRET-off, producing an array that indicated when FRET events started and ended. All reported FRET efficiencies (E_FRET_) were calculating by E_FRET_ = I_A_ / (I_A_ + I_D_), where I_A_ and I_D_ are the acceptor and donor intensities during a FRET-on event. The array of assigned states was processed with custom scripts to provide separate arrays for each indicated step in the single-molecule reaction, providing time-lengths for each step for all molecules analyzed. These time-length arrays were used to calculate cumulative probabilities, plotted, and fit to an exponential distribution function using cdfcalc on MATLAB. Association rates and dissociation rates (inverse lifetimes) were extracted from the exponential fit. The exponential equation used was:

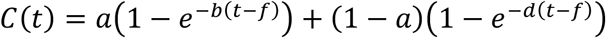

where t is time, a is the amplitude of molecules in a specific phase, b and d are rates for each phase, and f is an adjustment factor. If a corresponding phase had an amplitude less than 0.1, an exponential fit composed of only a single phase was used instead.

For the 43S PIC loading assays, data were processed on MATLAB 2024a as previously described. The cumulative probabilities for conditions showcasing two-step sequential pathways were fit to a hypoexponential distribution function defined as:

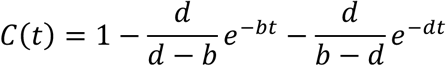

where t is time and b and d were the rates for each step. Reported errors for derived rates represent the 95% confidence interval produced from fits to linear, single-exponential, double-exponential, or hypoexponential functions as indicated. The reported mean times were defined as the reciprocal of the derived rate constant or weighted inverse rate constant in the case of double-exponential fits.

To produce fluorescence intensity heat maps for the PIC loading and mRNA extension assay, the data were normalized by computing a maximum and minimum for each fluorescent signal (Cy3.5-40S and Cy5-probe). For the Cy5-probe, maximum and minimum were determined from the top and bottom 10-15^th^ percentiles of intensity respectively. A similar approach was taken for Cy3.5-40S intensities but normalization was done only on data post mRNA extension, hence the negative normalized intensities observed on the plot prior to extension caused by fluorescent cross talk between Cy3 and Cy3.5.

To calculate errors in the efficiency of binding events or percent of molecules in FRET, bootstrap analyses (n = 10,000) were performed as described previously^43^.

## Acknowledgements

We are grateful to members of the Puglisi and Fraser labs for feedback and guidance. We thank Zev Bryant and Gheorghe Chistol for their feedback, discussions and guidance; Peter Sarnow and his lab for sharing cell culture equipment; and Baldandorj Baatar and Pierre-Jean Mattei for assistance with insect cell cultures. C.A. was supported by the Howard Hughes Medical Institute Gilliam Fellows Program. C.A. and C.I.S were supported by the Stanford Bio-X fellowship. This work was funded by a Chan Zuckerberg Biohub Investigator Award and the National Institutes of Health (GM145306 and AG064690) to J.D.P. C.P.L was supported by the NIH (GM160398). J.W. was supported by The Welch Foundation (I-2218-20240404). M.S. and C.S.F were supported by the NIH (GM152137 to C.S.F.).

## Author Contributions

Conceptualization, C.A. and J.D.P.; Methodology, C.A., C.P.L., J.W., M.S., R.G., A.A.D., C.I.S., M.Z.P., A.M., J.J., C.S.F., and J.D.P.; Resources, C.A., C.P.L., J.W., M.S., R.G., A.A.D., C.I.S., M.Z.P., A.M., J.J., C.S.F., and J.D.P.; Investigation, C.A., A.A.D., and C.I.S.; Visualization, C.A., C.P.L., A.A.D., and M.Z.P.; Funding acquisition, J.J., C.S.F., J.D.P.; Project administration, C.S.F., and J.D.P.; Supervision: C.P.L., J.W., C.S.F., and J.D.P.; Writing – original draft: C.A.; Writing – review & editing: C.A., C.P.L., J.W., M.S., R.G., A.A.D., C.I.S., M.Z.P., A.M., J.J., C.S.F., and J.D.P.

## Supplementary Figures

**Figure S1.**
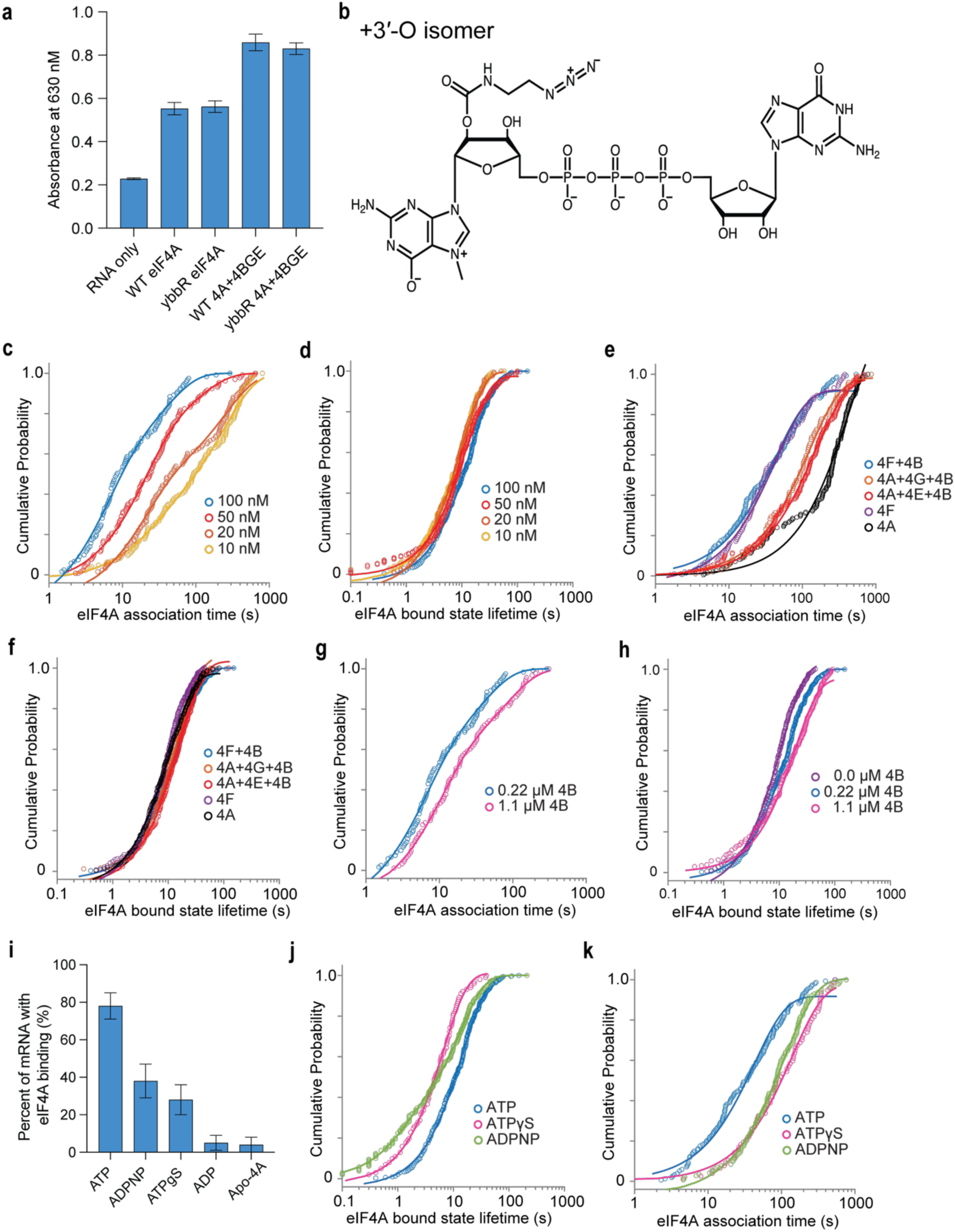
A 5′ end-to-eIF4A FRET assay. **a.** Bar plot of absorbance at 630 nm for the indicated conditions from a malachite green ATPase assay. Error bars represent standard deviation of data points, n = 2 for all conditions. **b.** Chemical structure of the cap analog (N_3_-m^7^GpppG) used for click chemistry labeling of capped mRNA. **c-h.** Cumulative probability plots of the indicated parameters at differing eIF4F concentrations reported by eIF4A concentration (**c** and **d**), differing eIF4 B, G, E dropouts (**e** and **f**), or differing 4B concentrations (**g** and **h**). Lines represent fits to exponential functions. Sample size is reported in Figures 1d, e. **i.** Bar plot of percent of mRNA molecules with Cy5-eIF4A binding events in differing ATP analog conditions. Error bars represent 95% CI from binomial bootstrapping. From left to right n = 111, 107, 121, 132, and 112. **j, k.** Cumulative probability plots of the indicated parameters with differing ATP analogs. Lines represent fits to exponential functions, sample size is reported in Figure 1d.

**Figure S2.**
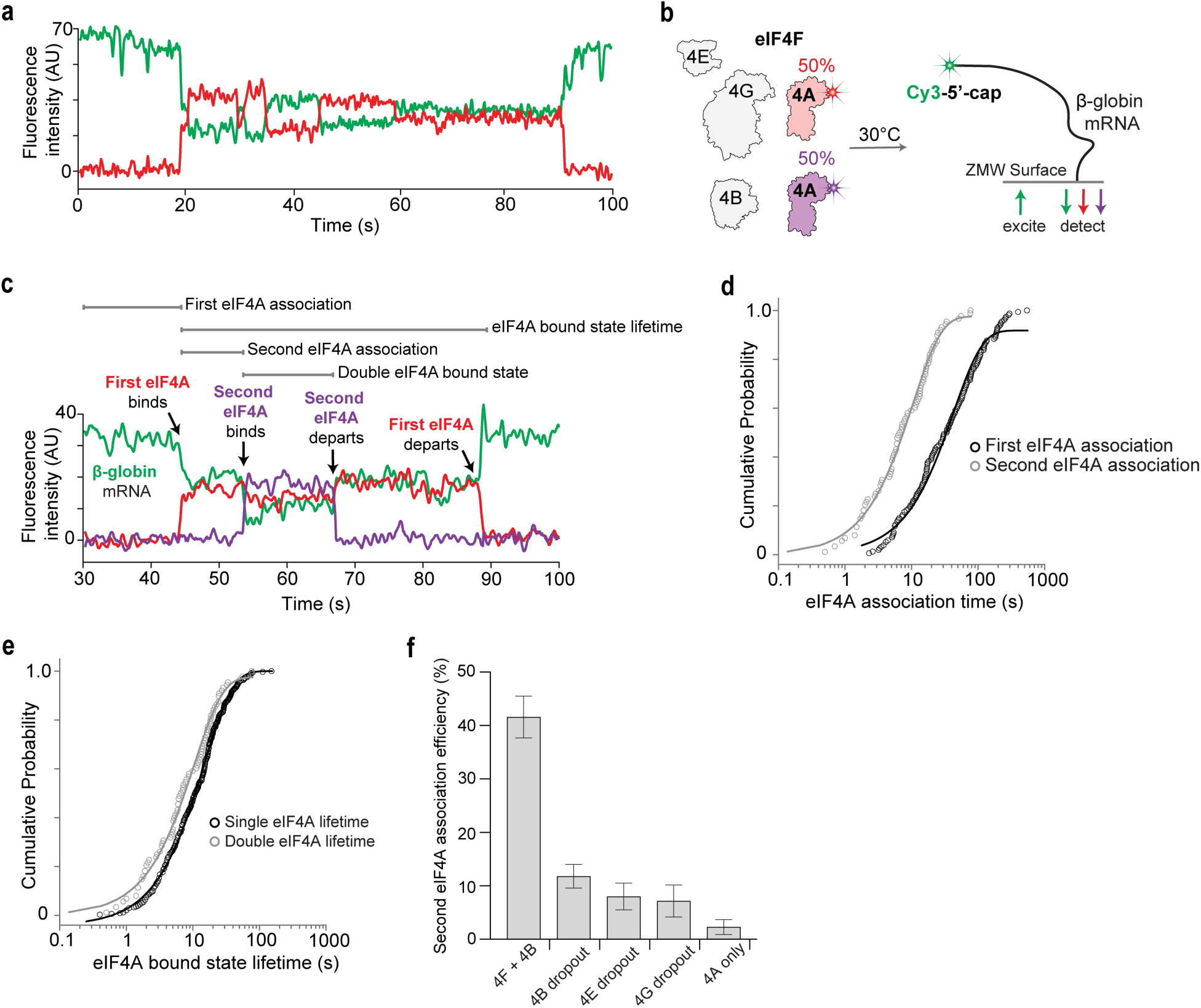
Multiple eIF4A bind proximal to the 5′ cap. **a.** Example single-molecule data showcasing multiple FRET states observed in the assay shown from Figure 1a. **b.** Schematic of single-molecule cap-to-eIF4A FRET assay where Cy5-labeled and Cy5.5-labeled eIF4A alongside eIF4E, eIF4G, eIF4B, and ATP are delivered to Cy3-5′-cap labeled β-globin mRNA immobilized on a ZMW imaging surface. **c.** Example single-molecule trace for the assay in (**b**), bursts in red or purple intensity anti-correlated with green intensity decreases (Cy3-labeled β-globin) indicate labeled eIF4A binding. The association times and lifetimes of multiple distinct eIF4A can be tracked. **d, e.** Cumulative probability plots of the indicated parameters, comparing first and second eIF4A binding events. The lines represent fits to exponential functions, for first binding n = 115, and for second binding n = 84. **f.** Bar plot for the fraction of mRNA with double eIF4A binding on the indicated conditions. From left to right n = 323, 177, 199, 201, and 210.

**Figure S3.**
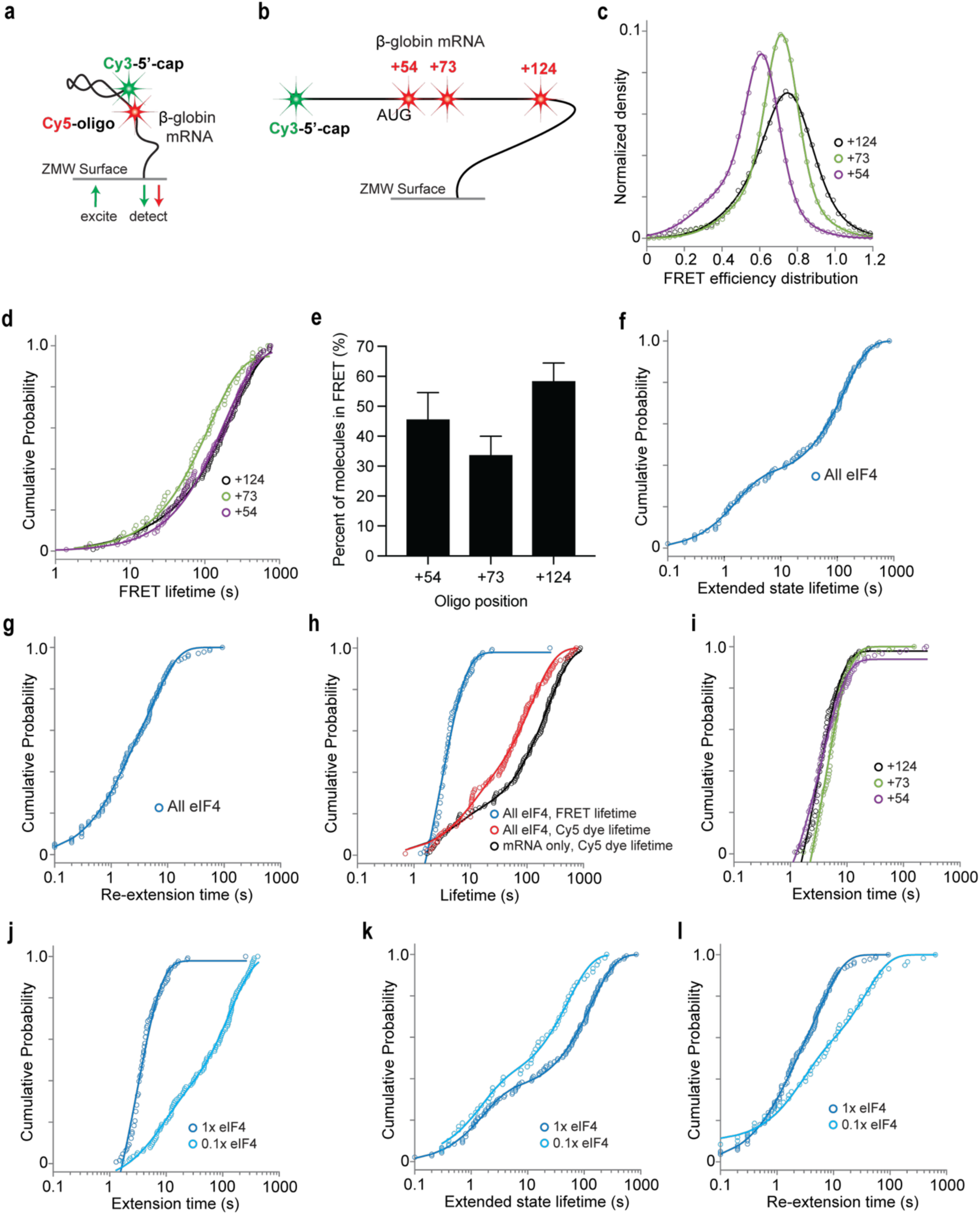
A cap-to-oligo FRET assay. **a.** Schematic of single-molecule assay to probe conformation at the 5′ end of β-globin mRNA. A Cy3-5′-cap labeled β-globin mRNA is hybridized to a Cy5-probe and immobilized on a ZMW imaging surface. **b.** Schematic showcasing the different Cy5- probe positions used relative to the 5′ cap. **c.** Plots of FRET efficiency distribution for the different probe positions. Lines indicate fits to Gaussian functions (Values for each variable in the fit displayed in Table S4). For +54, +73, and +124: n = 124, 56, and 121. **d.** Cumulative probability plots of the FRET lifetime for the indicated probe positions. Lines represent fits to exponential functions. **e.** Plots of percent of molecules in FRET at each probe position indicated. Error bars represent 95% CI from binomial bootstrapping. From left to right, n = 108, 200, 225. **f, g.** Cumulative probability plots of the indicated parameters with all eIF4 protein in cap-to-probe FRET assay using probe position +124. The lines represent fits to exponential functions. **h.** Cumulative probability plot of lifetimes for the indicated conditions. Lines represent fits to exponential functions. **i.** Cumulative probability plots of extension time comparing different Cy5-probe positions. Lines represent fits to exponential functions. For +54, +73, +124: n = 118, 130, and 131. **j-l.** Cumulative probability plots of the indicated parameters, at two different eIF4 protein concentrations. The lines represent fits to exponential functions, for 0.1x eIF4 n = 116 molecules were analyzed.

**Figure S4.**
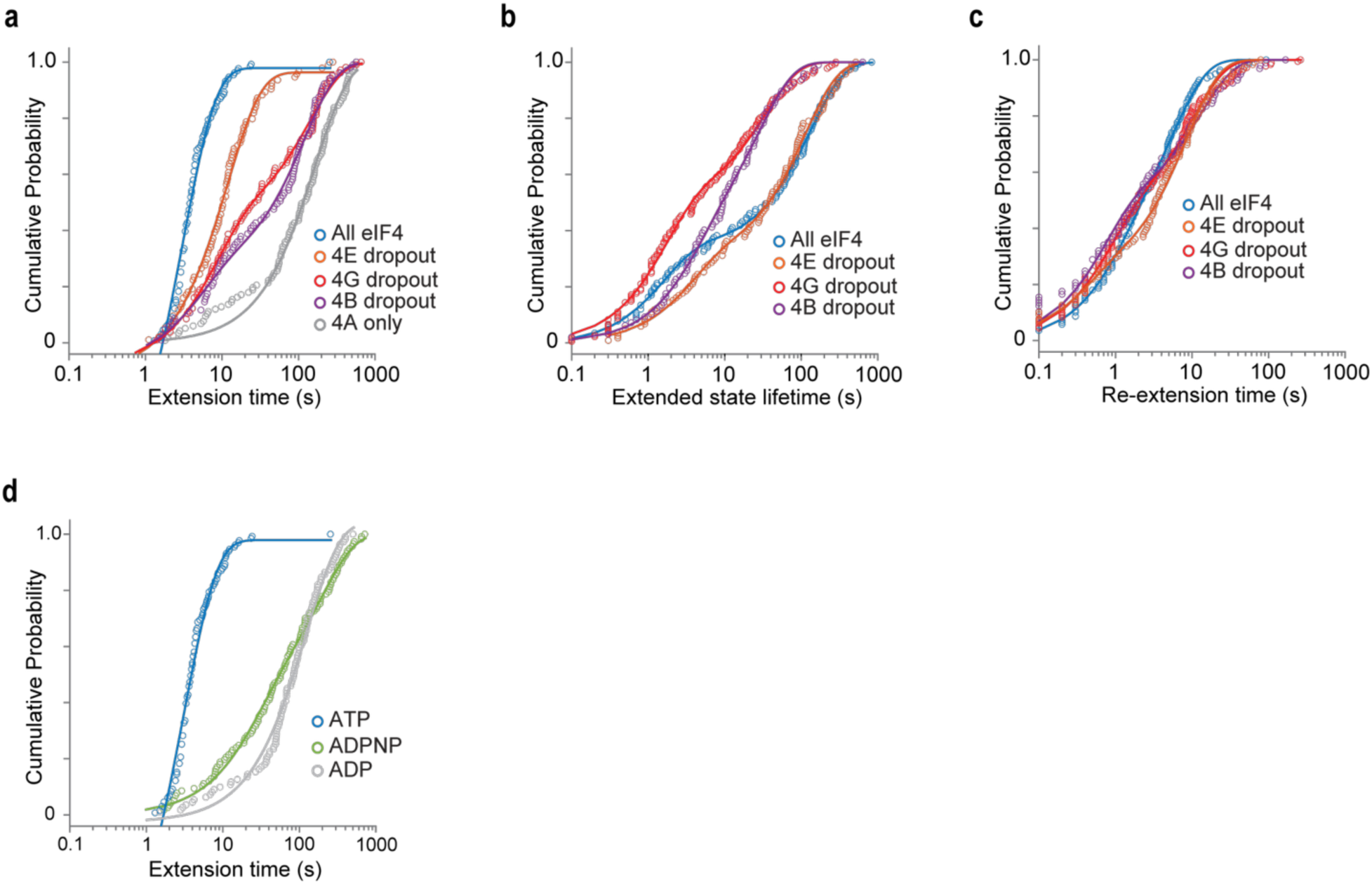
Factor omissions and ATP modulate mRNA extension a-d. Cumulative probability plots of the indicated parameters for either various eIF4B, 4G, or 4E dropouts (**a**-**c**) or different ATP analogs (**d**). The lines represent fits to exponential functions; sample size is detailed in Figure 3d.

**Figure S5.**
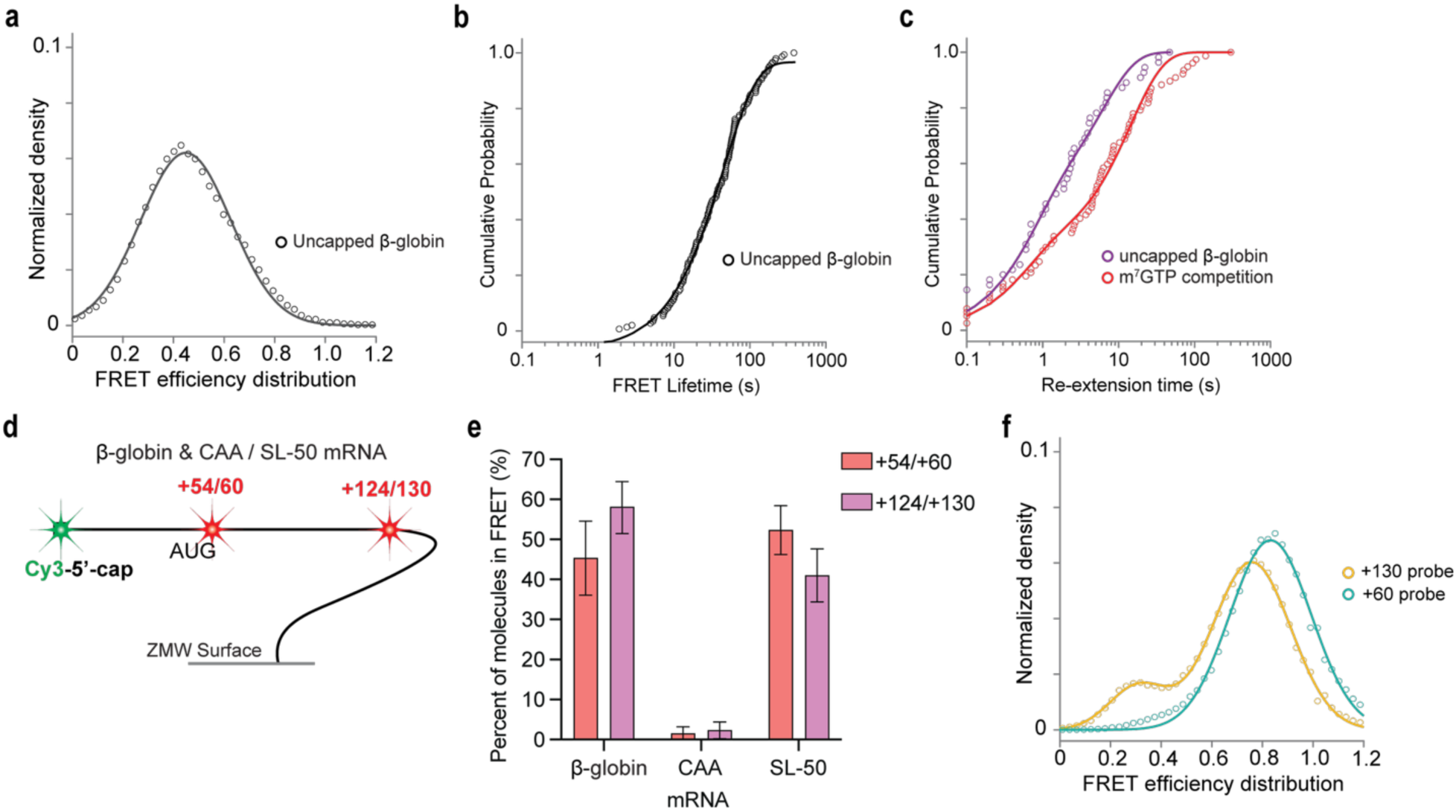
Characterizing model mRNA constructs in cap-to-oligo FRET assay. **a, b.** Plot of FRET efficiency distribution (**a**) and cumulative probability plot (**b**) for Cy3-labeled uncapped β-globin mRNA to Cy5-probe annealed at position +124. Line represents Gaussian function fit (**a**) (see Table S4 for values of the fit) or exponential function fit (**b**), n = 147. **c.** Cumulative probability plots of subsequent FRET lifetimes for the indicated conditions. Lines represent fits to exponential functions. **d.** Schematic showcasing the different Cy5-probe positions used relative to the 5′ cap with positions given for β-globin or CAA mRNA (+54 and +124) and SL-50 mRNA (+60 and +130). **e.** Bar plots of percent of molecules in FRET for the indicated mRNA at two different probe positions as shown in (**d**). Error bars represent 95% CI from binomial bootstrapping, from left to right n = 108, 225, 250, 251, 250, and 248. **f.** Plot of FRET efficiency distribution for Cy3-labeled SL-50 mRNA to Cy5-probe annealed at indicated positions. Sample size is reported in Figures 4e, f.

**Figure S6.**
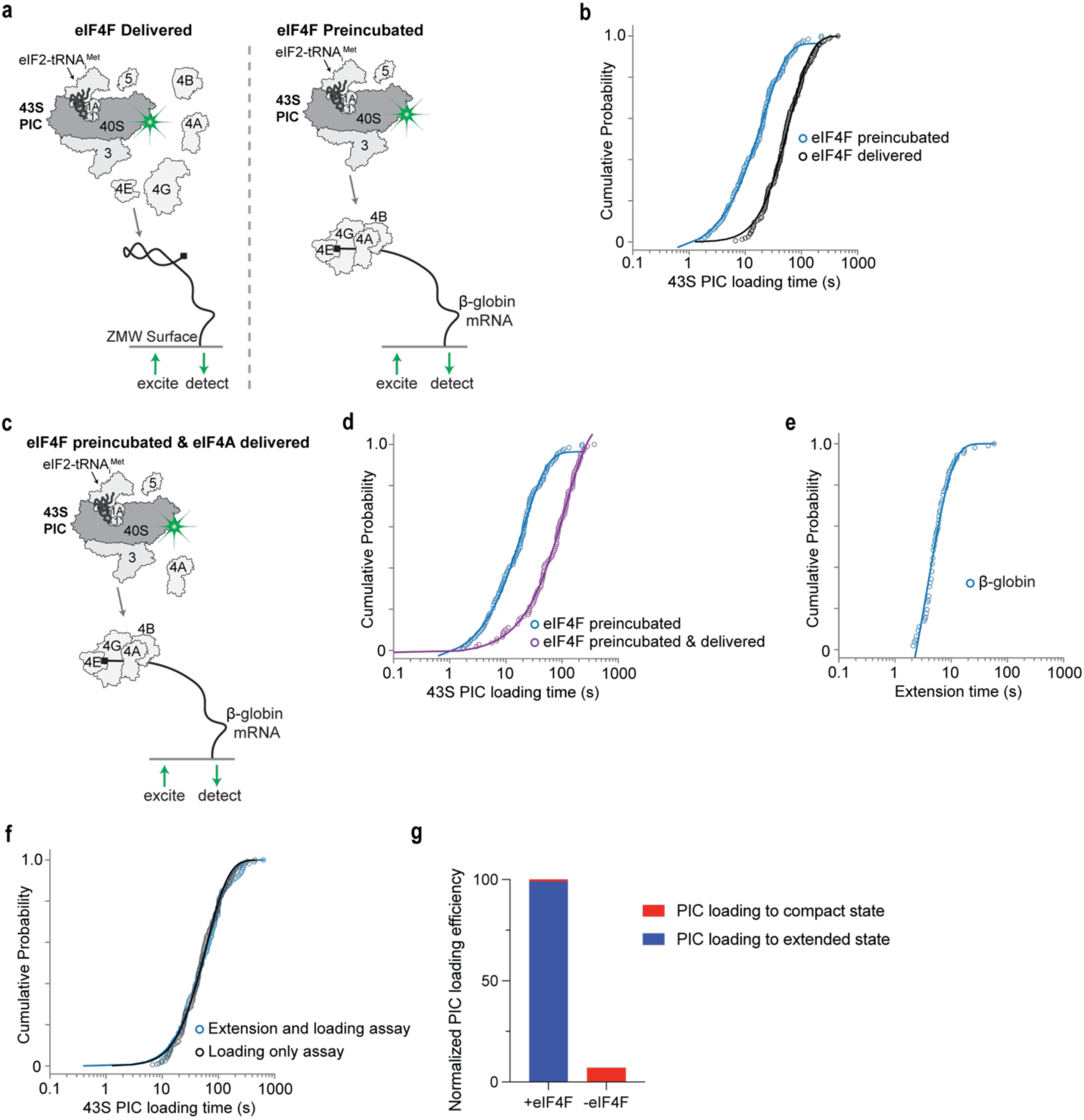
Extended data associated with Figure 5. **a.** Schematic denoting the different eIF4F conditions for the single-molecule 43S PIC loading assay. Delivered where all eIF4 proteins are delivered alongside the PIC or preincubated where the eIF4 proteins are added to the mRNA on the imaging surface prior to PIC delivery. **b.** Cumulative probability plots of 43S PIC loading time for the two conditions detailed in (**a**). Lines represent fits to either exponential (preincubated) or hypoexponential (delivered) functions; for delivered n = 156, for preincubated n = 132. **c.** Schematic showcasing the eIF4A preincubated and delivered condition for the single-molecule 43S PIC loading assay. **d.** Cumulative probability plots of 43S PIC loading time for the conditions indicated. Lines represent fits to exponential functions; for preincubated and delivered n = 109. **e.** Cumulative probability plot of extension time from assay tracking extension and PIC loading (Fig. 5d). Lines represent fits to exponential functions, n = 129. **f.** Cumulative probability plots of PIC loading times post-synchronized to delivery of the PIC comparing two separate assays (Fig. 5a and 5d). Lines represent fits to exponential functions. **g.** Plot of normalized PIC loading efficiency (fraction of mRNA with an observed 40S binding event) with and without eIF4F, blue and red shading denote the state of the mRNA when a PIC is loaded. From left to right, n = 168 and n = 164.

**Figure S7.**
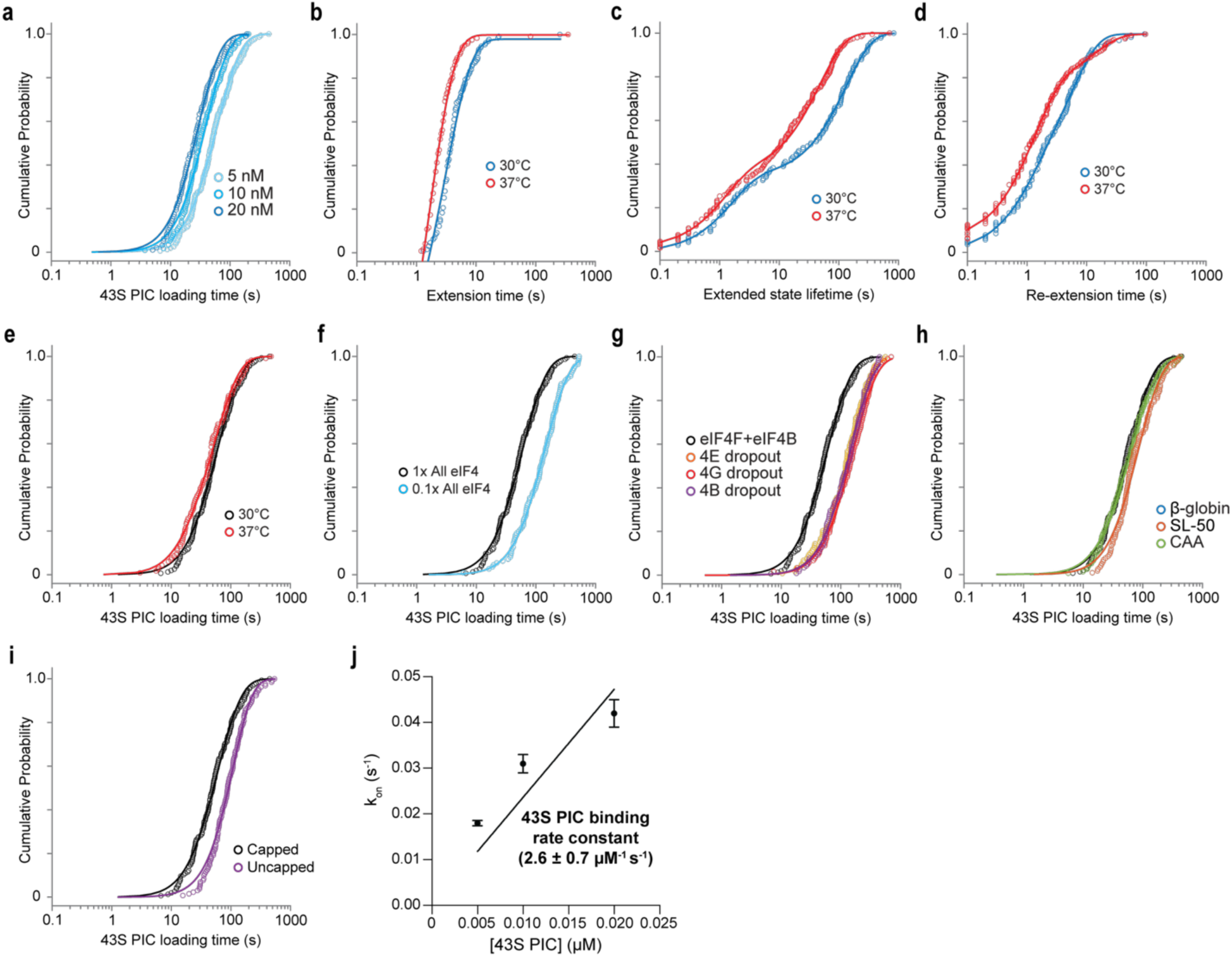
Cumulative probability plots associated with Figure 6. **a.** Cumulative probability plots of 43S PIC loading times at differing PIC concentrations. Lines represent fits to hypoexponential functions. Sample sizes displayed in Figure 6a. **b-e.** Cumulative probability plots of indicated parameters comparing two different temperatures of the reaction. Lines represent fits to either exponential (**b**-**d**) with n = 121 for 37 C condition, or hypoexponential functions (**e**) with n = 116 for 37 C condition. **f-i.** Cumulative probability plots of the 43S PIC loading at differing eIF4F concentrations reported by eIF4A concentration (**f**), differing eIF4 B, G, E dropouts (**g**), differing mRNAs (**h**), or differing capped states for β-globin (**i**). Lines represent fits to hypoexponential functions (**e**). In (**e**), the 0.1x eIF4 condition, n = 111. All other sample sizes were reported in Figures 6b-d. **j.** Plot of observed slow step in 43S PIC binding rate across different concentrations. Linear regression analysis (solid line) was used to derive the association rate constant. The error bars represent the 95% confidence interval (CI) of the observed association rate constants.

---

**Table S1. Summary of 5’ end-to-eIF4A FRET assay data**

All rates, rate constants, mean times, number of molecules, and number of binding events analyzed in each single-molecule experiment.

**Table S2. Summary of cap-to-probe FRET assay data**

All rates, rate constants, mean times, and number of molecules analyzed in each single-molecule experiment. Probe positions are listed relative to the 5’ cap.

**Table S3. Summary of 43S PIC loading assay data**

All rates, mean times, and number of molecules analyzed in each single-molecule experiment.

**Table S4. Summary of FRET efficiency distribution for cap-to-probe FRET assay** Values from Gaussian fits of FRET efficiency distribution in the cap-to-probe FRET assay. Fits are for conditions with mRNA only. Mean values (µ), standard deviations (σ), normalized amplitudes (a), and 95% CI for all variables are reported.

**Table S5.**
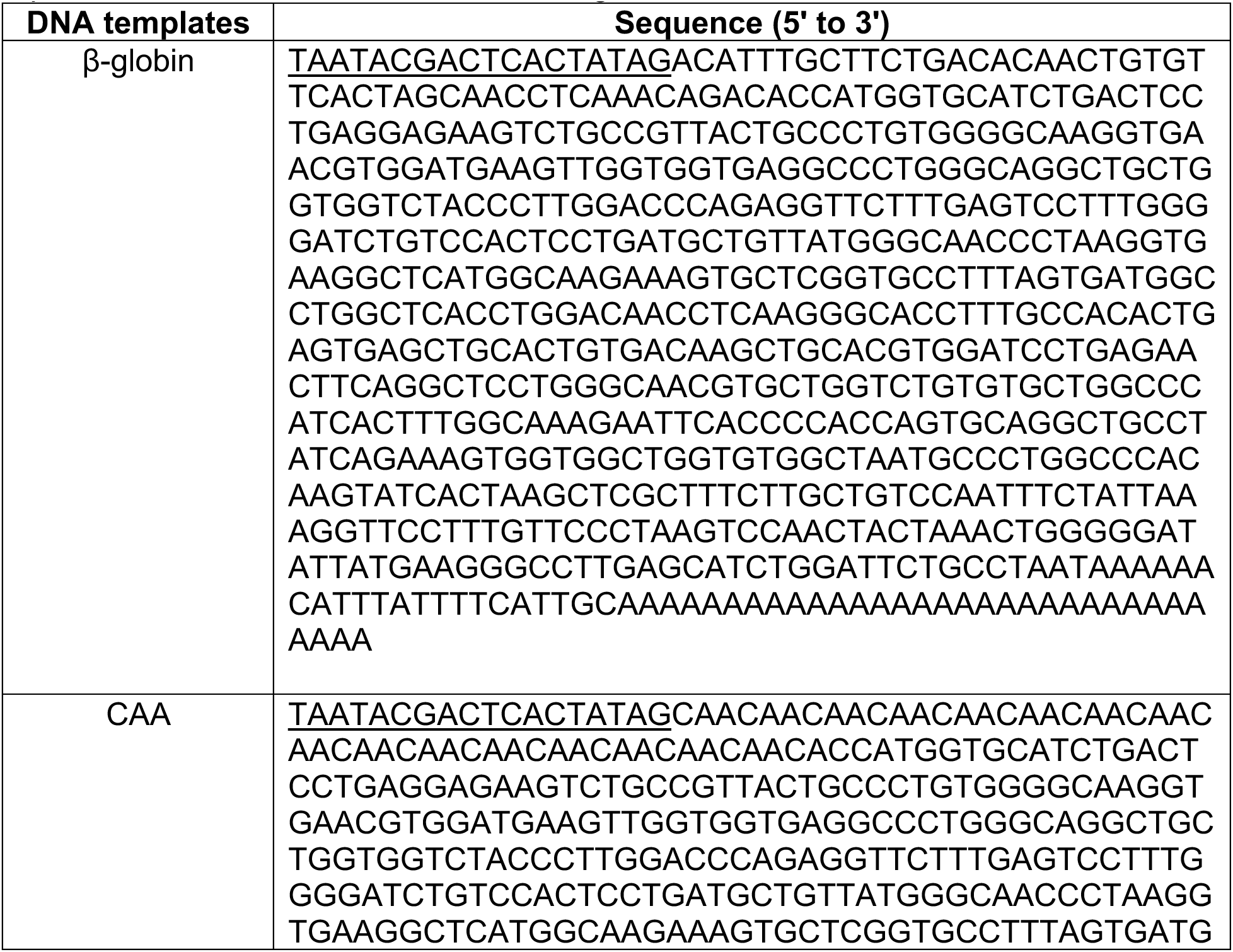
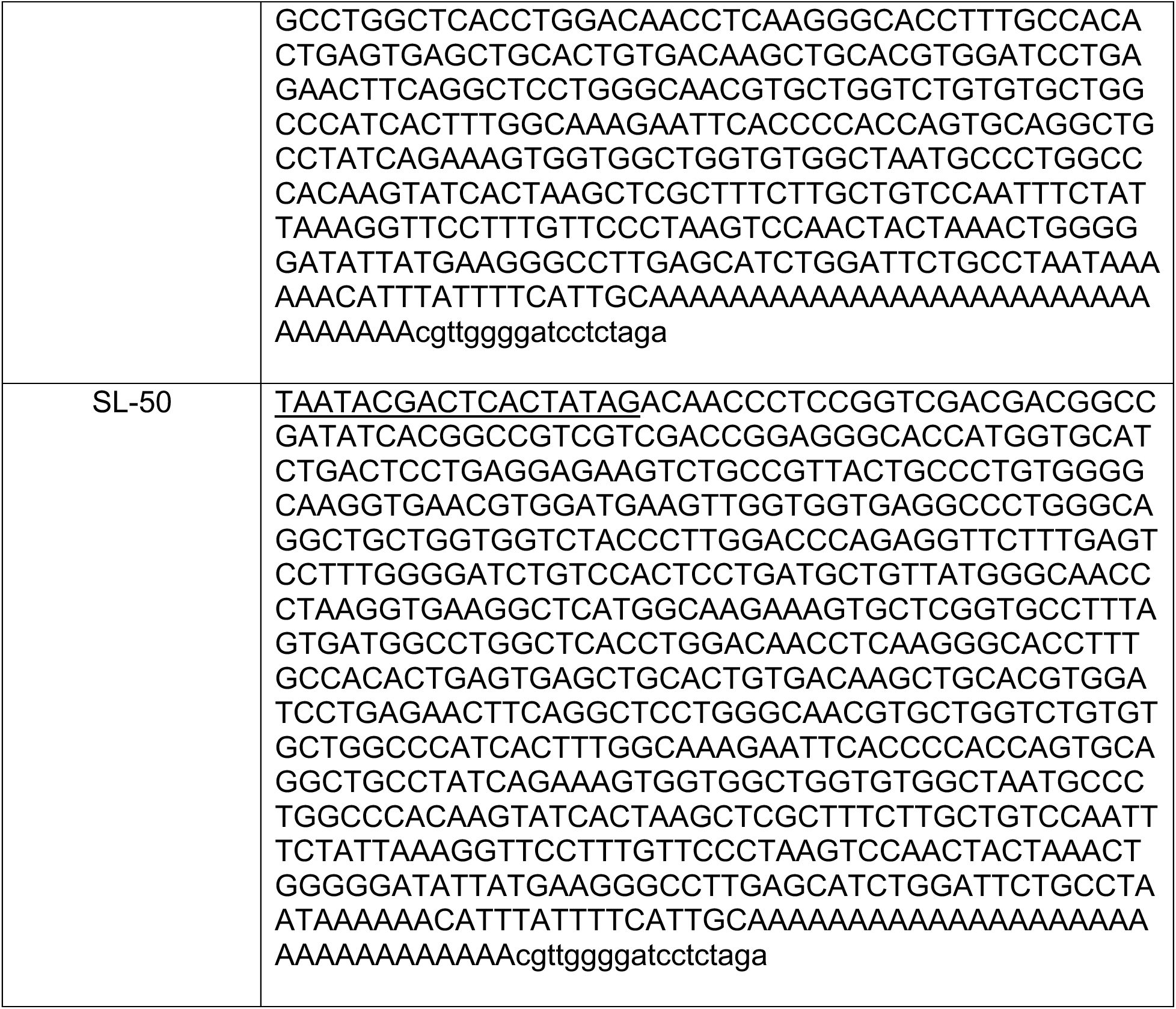
List of DNA templates used for mRNA *in vitro* transcription. Underlined section represents the T7 promoter sequence. Lower case sequence represents additional nucleotides remaining after linearization with XbaI.

**Table S6.**
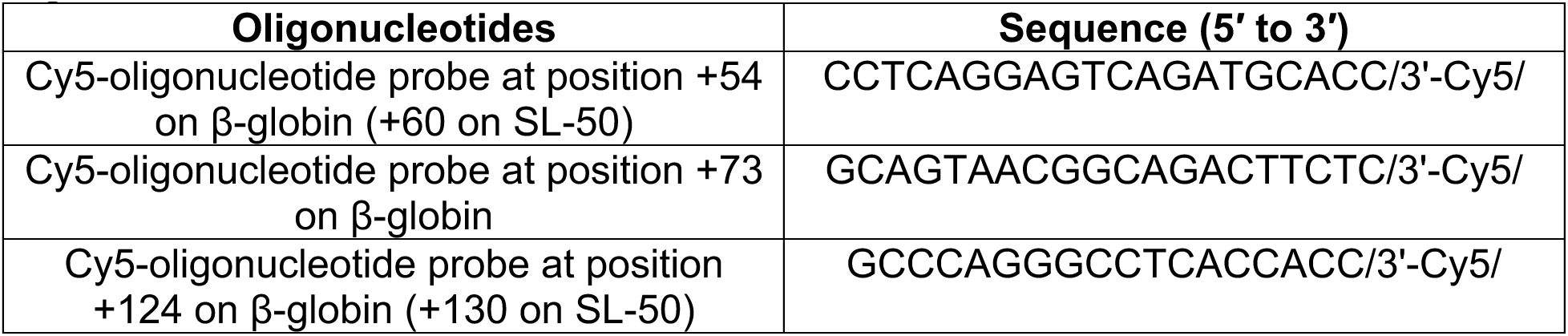
List of DNA oligonucleotide probes used in cap-to-probe FRET assay. The notation /3’-Cy5/ represents a Cy5 dye covalently tethered to the 3’ end of the oligonucleotide.

## Notes

### Competing Interest Statement

The authors have declared no competing interest.

